# Systematic comparison of monoclonal versus polyclonal antibodies for mapping histone modifications by ChIP-seq

**DOI:** 10.1101/054387

**Authors:** Michele Busby, Catherine Xue, Catherine Li, Yossi Farjoun, Elizabeth Gienger, Ido Yofe, Adrianne Gladden, Charles B. Epstein, Evan M. Cornett, Scott B. Rothbart, Chad Nusbaum, Alon Goren

**Author notes:** Correspondence should be addressed to Alon Goren. Email Addresses: Michele Busby, Catherine Xue, Catherine Li, Yossi Farjoun, Elizabeth Gienger, Ido Yofe, Adrianne Gladden, Charles B. Epstein, Evan M. Cornett, Scott B. Rothbart, Chad Nusbaum, Alon Goren.

## Abstract

**Background:** The robustness of ChIP-seq datasets is highly dependent upon the antibodies used. Currently, polyclonal antibodies are the standard despite several limitations: they are non-renewable, vary in performance between lots, and need to be validated with each new lot. In contrast, monoclonal antibody lots are renewable and provide consistent performance. To increase ChIP-seq standardization, we investigated whether monoclonal antibodies could replace polyclonal antibodies. We compared monoclonal antibodies that target five key histone modifications (H3K4me1, H3K4me3, H3K9me3, H3K27ac and H3K27me3) to their polyclonal counterparts in both human and mouse cells.

**Results:** Overall performance was highly similar for four monoclonal/polyclonal pairs, including when we used two distinct lots of the same monoclonal antibody. In contrast, the binding patterns for H3K27ac differed substantially between polyclonal and monoclonal antibodies. However, this was most likely due to the distinct immunogen used rather than the clonality of the antibody.

**Conclusions:** Altogether, we found that monoclonal antibodies as a class perform as well as polyclonal antibodies for the detection of histone post-translational modifications in both human and mouse. Accordingly, we recommend the use of monoclonal antibodies in ChIP-seq experiments.

## Background

Chromatin immunoprecipitation followed by sequencing (ChIP-seq) is one of the key technologies for investigating the genomic localization of DNA-associated proteins. The ChIP-seq approach can be performed in two major ways: native ChIP (where the original genomic localization of DNA associated proteins is maintained without cross-linking) and cross-linked ChIP. Here, we focused on the cross-linked ChIP-seq approach, as most of the public datasets relevant to our samples were produced by this method. In this technique, the DNA-associated proteins are cross-linked to the DNA. After DNA shearing, a specific antibody is used to enrich the targeted protein by immunoprecipitation, which also enriches the specific DNA it is bound to because it is cross-linked to it. Finally, the DNA fragments that precipitated with the enriched protein are sequenced. Hence, the results of each experiment are highly dependent upon the quality of the antibody that is used.

Polyclonal antibodies have been used as the standard antibody reagent for ChIP-seq by many laboratories and consortia [1–3]. Problematically, however, each polyclonal antibody lot is a limited resource, as each is raised from a different immunized animal. Each polyclonal antibody batch consists of a highly complex population of individual antibody molecules, representing the unique response of the source animal’s immune system. Some of these component antibody molecules will specifically target the epitope in question, but other molecules in this population may enrich for other off-target epitopes. Different antibody lots raised to the same target epitope will thus naturally differ in performance [4, 5] and each must be validated before use. Critically, once exhausted, a polyclonal antibody lot cannot be reproduced [6].

To overcome these limitations, many scientists have advocated for the use of monoclonal antibodies [7–9]. Monoclonal antibodies are harvested from purified cell lines derived from a single immune cell, which brings distinct advantages: first, lots consist of a single antibody species that specifically targets the desired epitope; second, monoclonal lots are uniform in performance; and third, lots are renewable resources as long as the cell line is maintained. Approaches that attempt to overcome the limitations of polyclonal antibodies include the development and optimization of recombinant antibodies [10], development of recombinant antibodies that provide “antigen clasping” [11], the generation of specific monoclonal antibodies followed by evaluation of their performance [12–14] and the comparison of multiple antibodies targeting repressive histone modifications [15].

However, despite the advantages of monoclonal antibodies and the progress toward other approaches, citation data aggregated in the CiteAB database [16] indicates that polyclonal antibodies are used in published research more frequently than monoclonal antibodies (54% of citations versus 46% [17]); Similarly, in a study conducted as part of the NIH modENCODE [18] and Roadmap Reference Epigenome [2] projects, about 74% (181 out of 246) of the histone modification antibodies surveyed were polyclonal [5].

To systematically investigate whether monoclonal antibodies can substitute for polyclonal antibodies in ChIP-seq procedures while retaining equivalent performance, we designed and carried out a direct side-by-side comparison. We compared a set of five monoclonal antibodies targeting key histone modifications (H3K4me1, H3K4me3, H3K9me3, H3K27ac, and H3K27me3) to their polyclonal counterparts, using the same antibodies and lots that had been previously validated by the ENCODE project [1] (**Table 1**). To ensure that all samples and antibodies were handled in a precisely controlled manner, all work was performed employing automated ChIP-seq protocols implemented on a standard laboratory liquid handling system.

**Table 1:** Antibodies used in the study.

| Epitope | Antibody Type | Commercial company | Catalog number | Lot IDs | Validation data |
| --- | --- | --- | --- | --- | --- |
| H3K4me1 | Monoclonal | CST (Cell Signaling Technology) | 5326 | 1,2 | https://www.encodeproject.org/antibodies/ENCAB650MWL/ |
| H3K4me1 | Polyclonal | Active Motif | 39297 | 1714002 | Supplemental Figure 7 |
| H3K4me3 | Monoclonal | CST | 9751 | 1,6,8,9 | https://www.encodeproject.org/antibodies/ENCAB902NZL/ |
| H3K4me3 | Polyclonal | Millipore | 17-614 | DAM1644057 | https://www.encodeproject.org/antibodies/ENCAB000BLE/ |
| H3K9me3 | Monoclonal | CST | 13969 | 2 |  |
| H3K9me3 | Polyclonal | Abcam | ab8898 | GR131093-3 | https://www.encodeproject.org/antibodies/ENCAB369JSU/ |
| H3K27ac | Monoclonal | CST | 8173 | 1,3 | https://www.encodeproject.org/antibodies/ENCAB502OHI/ |
| H3K27ac | Monoclonal | Active Motif | 39685 | Lot 35813005 | http://www.histoneantibodies.com/FinalArrayData/H3K27ac/ |
| H3K27ac | Polyclonal | Active Motif | 39133 | 31610003 | https://www.encodeproject.org/antibodies/ENCAB000AQN/ |
| H3K27me3 | Monoclonal | CST | 9733 | 8,10 | https://www.encodeproject.org/antibodies/ENCAB155VEG/ |
| H3K27me3 | Polyclonal | Millipore | 07-449 | 2064519 | https://www.encodeproject.org/antibodies/ENCAB036YAO/ |

As a class, we found that the performance of monoclonal antibodies targeting histone post-translational modifications in ChIP-seq assays matched the performance of polyclonal antibodies. Given that monoclonal antibodies represent a renewable resource, and eliminate the lot-to-lot variability that is expected with polyclonal antibodies, the replacement of polyclonal antibodies with monoclonal antibodies for use in ChIP-seq and similar affinity-based methods has significant benefits. Employing monoclonal antibodies will result in increased reproducibility and robustness and will substantially improve standardization of results among data sets.

## Results

We designed an experimental system for rigorously comparing the performance of monoclonal and polyclonal antibodies in ChIP-seq and applied it to antibodies targeting five key histone modifications (H3K4me1, H3K4me3, H3K9me3, H3K27ac and H3K27me3) (**Table 1**). These epitopes provide a rigorous test set of antibodies as they represent open and closed chromatin environments, have distinct localization patterns as described in **Table 2**, and are commonly used in studies of genomic organization of DNA associated proteins. We performed ChIP-seq with these antibodies in the human erythroleukemic cell line K562, the human lymphoblastoid cell line GM12878 and mouse embryonic stem (mES) cells. To control for experimental variability, we implemented a fully automated ChIP-seq process [19] that ensures precise liquid handling, maximizes reproducibility, and controls for human error. We performed two to four technical replicates for each antibody tested to control for experimental variability and sequenced the libraries using Illumina paired-end reads. To provide evidence for consistency between monoclonal lots, in a subset of the samples, we repeated the ChIP-seq with a distinct antibody lot. We then further computationally normalized our datasets to account for possible technical variability introduced by fragmentation and differing read depths. Finally, we analyzed our data to compare the performance of monoclonal and polyclonal antibodies focusing on the specificity and the number of peaks identified, as well as the overall pattern of reads localized across the genome.

**Table 2:** Datasets summary.

| Antibody | Number of Replicates |  | Region targeted |
| --- | --- | --- | --- |
|  | Mono | Poly |  |
| H3K27ac | 6<br>(2 in K562;<br>2 in GM12878<br>and 2 in mES) | 2 (in K562) | Transcription start<br>sites, enhancers |
| H3K27me3 | 8<br>(4 in K562; 2<br>in GM12878;<br>2 in mES) | 3 (in K562) | Repressed regions |
| H3K4me1 | 7<br>(3 in K562; 2<br>in GM12878;<br>2 in mES) | 2 (in K562) | Enhancers |
| H3K4me3 | 14<br>(6 in K562; 4<br>in GM12878;<br>4 in mES) | 4 (in K562) | Transcription start<br>sites |
| H3K9me3 | 7<br>(3 in K562; 2<br>in GM12878;<br>2 in mES) | 2 (in K562) | Repressed regions |

### Normalization of ChIP-seq datasets

Before analyzing our data, we computationally normalized the aligned reads to isolate the effects of each antibody from two possible issues that could confound the comparison: (**i**) a higher number of reads increases the power to distinguish peaks from background noise [20]; (**ii**) chromatin DNA has been shown to shear into different size fragments in regions of open versus closed chromatin, and genomic regions originating from open chromatin are more likely to shear into small fragments [21]. The combination of this shearing bias and a narrow size selection can lead to an artifactual enrichment of reads in areas of open chromatin leading to pile ups of reads that mimic peaks.

The effects of fragment length bias are therefore protocol-specific and dependent upon both the fragmentation method and size selection. To quantify the effect of fragmentation on the localization of reads in our protocol, we examined our WCE control data. First, we defined the regions as open or closed chromatin based on ENCODE mappings derived from the combined annotations of ChromHMM [22] and Segway [23]. This mapping approach annotates the K562 genome according to seven canonical types: transcription start sites, promoter flanking regions, enhancers, weak enhancers, CTCF-enriched elements, transcribed regions and repressed regions. According to this mapping approach, the majority of the annotated K562 genome (84%) is in repressed regions (closed chromatin) while only ~1% of the K562 annotated genome is in transcription start sites (open chromatin).

Next, to assess the regional bias of the fragmentation of the cross-linked DNA, we quantified insert sizes of fragments falling into open and closed chromatin, expecting the insert size to be equivalent to the size of the DNA fragment originating in the immunoprecipitation step. To explore the effects of fragment length variation in our system, we examined reads with insert sizes between 70 and 700 bases, the size range of inserts typically found on an Illumina flow cell. We observed that the percentage of reads localizing to transcription start sites (TSS) was inversely correlated with the length of the insert size (R^2^=0.80) with a 2.6-fold higher percentage of reads localizing to TSS in read pairs with shorter (70–120bp) versus longer (650–700bp) insert sizes. Reads localizing to repressed regions were positively correlated with insert size (R^2^=0.70) though the difference in coverage is only 5% (**Supplemental Figure 1**).

While we have optimized our shearing process to provide high reproducibility of the fragmentation process (**Methods**), we recognized the possibility that fragmentation performance could vary among samples since each sample is sheared independently. To account for potential differences in fragmentation, we randomly selected alignments so that each aligned read set for a given histone modification had the same number of reads and fragment size distribution (**Supplemental Table 1**). As the insert size is equal to the length of the DNA fragment in the original pool, this normalization method approximates experiments that have both the same fragmentation and read depth.

### Comparison of peaks between ChIP-seq datasets

We investigated the relative performance of the antibodies in terms of sensitivity, specificity, and the number and distribution of peaks. Initial visualization of the data in a genome browser revealed a high degree of similarity in read coverage between monoclonal and polyclonal antibodies (**Figure 1 and Supplemental Figure 2**).

**Figure 1.**
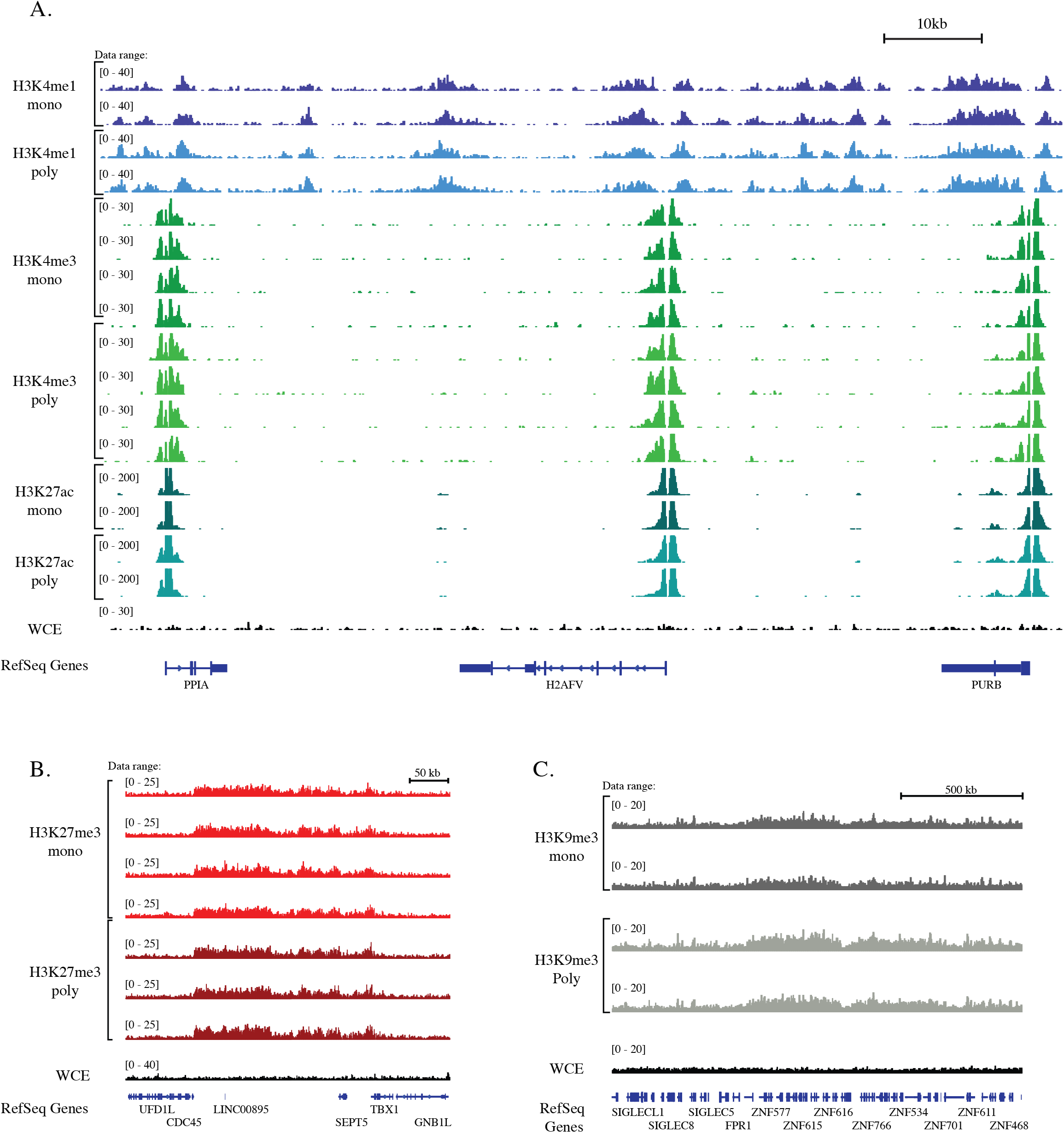
Read coverage across the genome. Images of Tiled Data Files (TDFs) generated by the IGV browser [34, 35] displaying the density tracks of reads aligned across the genome. The tracks show the correspondence in read coverage in monoclonal and polyclonal antibodies over representative genomic loci. **A**. Chromosome 7: 44,829,782-44,930,648 (about 100Kb), shows the read coverage of histone modifications associated with ‘active chromatin’ (H3K4me1, H3K4me3 and H3K27ac). The correspondence of read coverage of datasets for two major histone modifications associated with repression: **B.** H3K27me3 (Chromosome 22:19,492,023-19,849,594 (about 350Kb)), and **C.** H3K9me3 (Chromosome 19: 51,746,058-53,362,194 (about 1.6Mb)).

The best performing antibodies in ChIP-seq are those that provide the highest enrichment for DNA fragments associated with the target protein. However, measuring antibody enrichment is challenged by the absence of a set of known genomic patterns for histone modifications to serve as a baseline. For example, a greater portion of reads localizing to observed peaks could be indicative of either higher sensitivity of the antibody for its epitope or the addition of false peaks resulting from a higher degree of non-specific binding. For this reason we evaluated antibody performance using several different approaches.

We first sought to compare the locations of the peaks called in data from each antibody type. The ability to call peaks is a function of both antibody specificity and read depth. Thus, the analysis of peak localization ideally requires the unambiguous localization of peaks. To control for replicate-specific variability and provide deeper read coverage, we merged the data from the technical replicates to create a larger set of reads. This deeper dataset allowed us to assess the depth of sequencing coverage beyond which additional sequencing would not improve peak call accuracy. We randomly downsampled each dataset to twenty different read depths and called peaks using the HOMER version 4.7 peak caller [24] on the down-sampled read sets, using the default parameters for histone marks. At each sequencing depth, we determined the number of bases of the genome that were identified as being in a peak. The number of genome bases identified as being in peaks increased and then reached saturation with increasing read depth for each of the H3K27ac, H3K4me1, and H3K4me3 datasets (**Figure 2A**) but did not appear to reach saturation for H3K27me3 or H3K9me3 (**Supplemental Figure 3**). Each of the antibodies, monoclonal and polyclonal, followed the same pattern of saturation as its counterpart, indicating that regardless of antibody type the datasets required approximately the same depth of sequencing and that the differences between them cannot be overcome by deeper sequencing.

**Figure 2.**
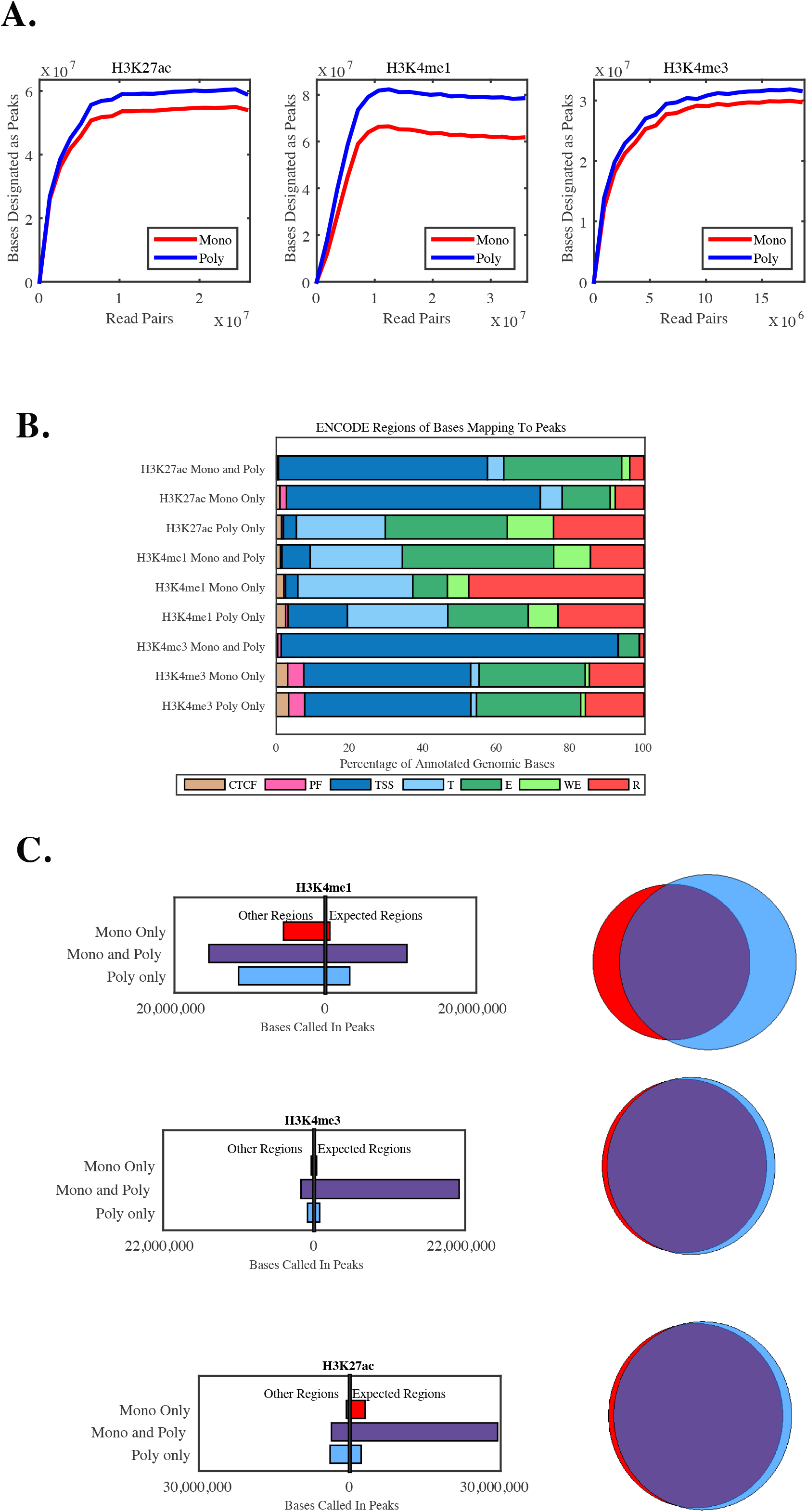
**A.** Saturation curve showing the number of bases called as being in peaks as a function of sequencing depth. The final dataset of the merged technical replicates was randomly downsampled to 20 different read depths and peaks were called in each dataset using HOMER. **B.** Distribution of the canonical ENCODE regions of the genomic bases identified as being in peaks. Note that distribution of bases called in both the monoclonal and polyclonal antibody differs from the distribution of bases called by only one antibody with fewer bases in their expected regions. **C. Left:** Bases of the genome that were designated as peaks were identified as being in the expected canonical ENCODE region versus other regions. Only genomic bases annotated in the ENCODE segmentation tracks for K562 are included in this calculation. **Right:** Venn diagrams displaying the overlap of peak calls in the monoclonal and polyclonal antibodies. The bases of the genome are identified as being in peaks by the monocolonal (red), polyclonal (blue), or both (purple) antibodies.

Next, we focused our analysis of peaks on the histone modifications associated with open chromatin (H3K27ac, H3K4me1, and H3K4me4), as in these datasets we were able to call peaks at a saturated read depth. For each of these histone modifications, more genomic bases were identified as being in peaks in the datasets for polyclonal antibodies than for monoclonal (**Table 3**). However, regions found in peaks for *both* types of datasets (**Figure 2**, Venn diagrams, **purple**) demonstrated a higher association with canonical ENCODE regions than ones that are found only in the polyclonal or only in the monoclonal datasets. Using the canonical ENCODE regions as a proxy for the true regions, we found that the polyclonal antibodies showed an increase in sensitivity at the expense of specificity. Nonetheless, the differences in both metrics were small, and data generated with both the monoclonal and polyclonal antibodies showed a high degree of consistency in determining which genomic bases were within peaks. Of the total genome bases that were identified by either antibody type as being in peaks, 77% (H3K27ac), 56% (H3K4me1), and 90% (H3K4me3) were identified by both types.

**Table 3:** Sensitivity and Specificity data for histone modifications associated with open chromatin. Sensitivity and specificity of monoclonal and polyclonal antibodies. Specificity is calculated as the percentage of the genomic bases that are identified as peaks that are within the expected canonical genomic region, as annotated by ENCODE. Sensitivity is calculated at the percentage of bases within the expected genomic region that are identified as being within peaks. Only genomic bases annotated in the ENCODE segmentation tracks for K562 are included in this calculation.

|  | Specificity |  | Sensitivity |  |
| --- | --- | --- | --- | --- |
|  | Mono | Poly | Mono | Poly |
| **H3K27ac** | **89%** | **85%** | **68%** | **69%** |
| **H3K4me1** | **38%** | **36%** | **54%** | **59%** |
| **H3K4me3** | **91%** | **90%** | **86%** | **87%** |

### Enrichment in Peaks

To further assess the specificity of binding, we used the peaks called in the merged datasets for each of the three antibodies associated with open chromatin to calculate a SPOT score [25] on each of the technical replicates. We found that the SPOT scores were slightly higher for the polyclonal antibody in H3K4me1 (p<0.01, average of 18% monoclonal versus 24% for the polyclonal) and in H3K4me3 (p<0.01, 27% monoclonal versus 32% polyclonal) but did not differ significantly for H3K27ac (p>0.05, 54% monoclonal versus 55% polyclonal). To assess the specificity in the marks associated with closed chromatin, we used the reference peaks called by ENCODE in K562 for H3K27me3 (ENCFF001SZF) and H3K9me3 (ENCFF001SZN) and calculated the percentage of reads in each dataset falling into these peaks. We found that in both cases the SPOT scores were nearly identical (36% and 38% in monoclonal (p<0.05 due to low variance) and polyclonal in H3K27me3 and 42% and 40% (p>0.05) in monoclonal and polyclonal in H3K27me3) indicating a high concurrence of read coverage.

### Specificity of binding

Next, we assessed all of the reads mapped to the genome to determine whether they were mapped to their expected regions. **Figure 3** and **Supplemental Figure 4** show the number of reads that mapped to each of the seven ENCODE canonical regions for each antibody. While results between the monoclonal and polyclonal antibodies for each epitope were similar, a greater percentage of reads mapped to their expected region of the genome (**Table 4**) for the polyclonal antibody to H3K4me3 (34% polyclonal mapping to transcription start sites vs 24% monoclonal, p<0.01). Due to the low variability between technical replicates in our system, small differences also reached statistical significance for the antibodies H3K27me3 (86% mono, 87% poly, p<0.05) and H3K9me3 (85% mono and 86% poly, p<0.05). We note that this approach – evaluating the percentage of reads mapped to ENCODE canonical genomic regions – does not provide a fully orthogonal validation of the specificity of the antibodies as the annotations were themselves created from ChIP-seq data.

**Figure 3.**
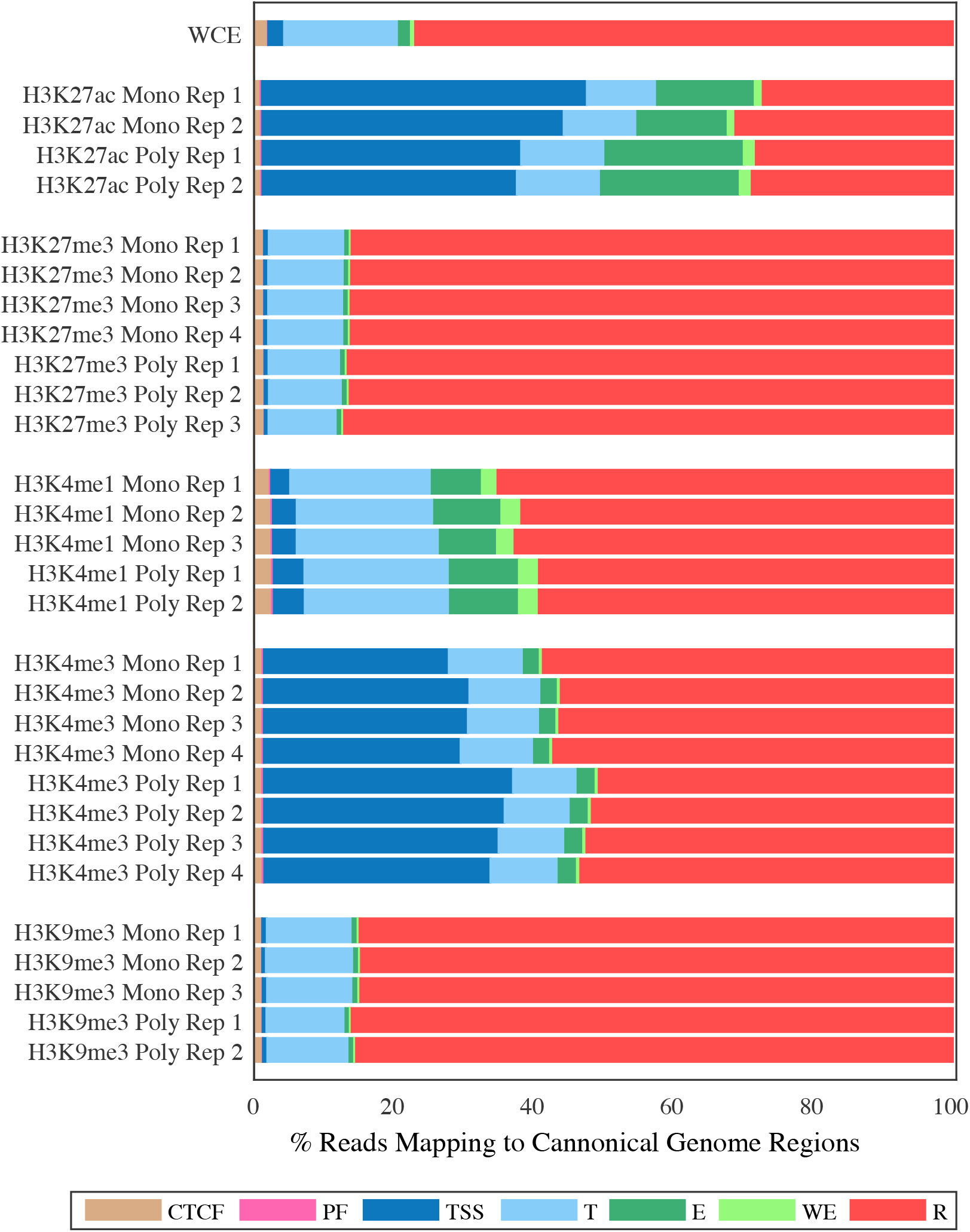
Reads in peaks mapping to canonical chromatin regions of the genome as defined by the ENCODE mappings. This plot displays the percentage of reads that map to each canonical genome region. The canonical genome regions were defined by the combined ENCODE mapping and are abbreviated as follows: CTCF-enriched elements (*CTCF*), promoter flanking regions (*PF*), transcription start sites (*TSS*), transcribed regions (*T*), enhancers (*E*), weak enhancers (*WE*), and repressed regions (*R*). Only reads that were located at regions identified as peaks were used for this plot. For each peak dataset the reads were normalized by insert size.

**Table 4:** Comparison of the percentage of reads in their expected E NCODE canonical regions (as defined in Table 2) between ChIP-seq datasets derived obtained by monoclonal and polyclonal antibodies. Enrichment versus WCE is defined as the percentage of reads in that region type in the sample divided by the percentage of reads in that region type in the WCE control.

|  |  | Percent Reads in Expected Regions | Enrichment Over WCE |
| --- | --- | --- | --- |
| H3K27ac | MonoRep1 | 60.5% | 15.2 |
|  | MonoRep2 | 56.1% | 14.1 |
|  | PolyRep1 | 56.8% | 14.2 |
|  | PolyRep2 | 56.3% | 14.1 |
| H3K27me3 | MonoRep1 | 86.1% | 1.1 |
|  | MonoRep2 | 86.1% | 1.1 |
|  | MonoRep3 | 86.2% | 1.1 |
|  | MonoRep4 | 86.2% | 1.1 |
|  | PolyRep1 | 86.6% | 1.1 |
|  | PolyRep2 | 86.3% | 1.1 |
|  | PolyRep3 | 87.1% | 1.1 |
| H3K4me1 | MonoRep1 | 7.2% | 4.2 |
|  | MonoRep2 | 9.6% | 5.6 |
|  | MonoRep3 | 8.2% | 4.8 |
|  | PolyRep1 | 9.9% | 5.8 |
|  | PolyRep2 | 9.9% | 5.7 |
| H3K4me3 | MonoRep1 | 26.5% | 11.7 |
|  | MonoRep2 | 29.4% | 13.0 |
|  | MonoRep3 | 29.2% | 12.9 |
|  | MonoRep4 | 28.2% | 12.4 |
|  | PolyRep1 | 35.6% | 15.7 |
|  | PolyRep2 | 34.4% | 15.2 |
|  | PolyRep3 | 33.6% | 14.8 |
|  | PolyRep4 | 32.4% | 14.3 |
| H3K9me3 | MonoRep1 | 84.9% | 1.1 |
|  | MonoRep2 | 84.7% | 1.1 |
|  | MonoRep3 | 84.8% | 1.1 |
|  | PolyRep1 | 86.0% | 1.1 |
|  | PolyRep2 | 85.4% | 1.1 |

### Whole Genome Read Coverage

We next investigated the distribution of ChIP-seq reads across the genome. To provide a basis for this quantitative evaluation, we defined non-overlapping bins of 2000 base pairs across the genome and counted the reads falling into each bin. We first compared the correlations in technical replicates in the samples normalized by insert size versus those normalized by random sampling. Correlations were highly similar, indicating that fragmentation and size selection were well controlled in these samples and did not introduce a significant source of bias (**Figure 4** and **Supplemental Figure 5**).

**Figure 4.**
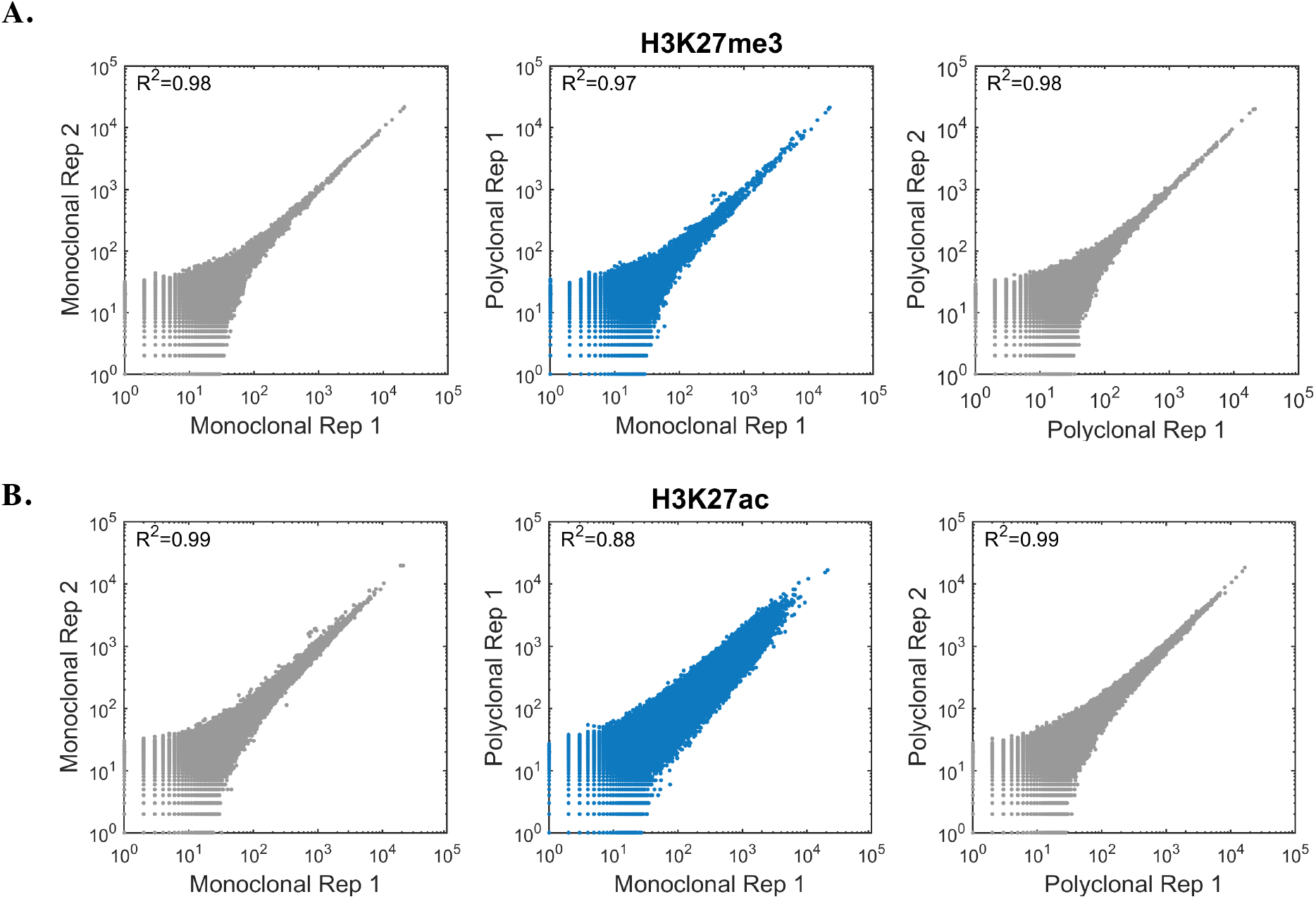
Correlation between monoclonal and polyclonal antibodies across the genome. Scatter plots (Loglog) presenting counts of reads per bin in non-overlapping 2000 bp windows tiled throughout the genome in replicates of the monoclonal antibody (**left**; gray), the polyclonal antibody (**right**; gray), and polyclonal versus monoclonal (**center**; blue). The H3K27ac data (**A**) show divergence between polyclonal and monoclonal antibodies, while the H3K27me3 data (**B**) show that the reproducibility is nearly indistinguishable from the reproducibility of data derived from technical replicates using the same antibody.

For all antibodies except H3K27ac the correlations between monoclonal and polyclonal antibodies were similar to those observed between technical replicates using the same antibody (**Figure 4** and **Supplemental Figure 5**).

Next, we examined the differences between the H3K27ac monoclonal and polyclonal samples more closely. The H3K27ac modification is present both at enhancer regions and transcription start site regions [26] so we compared the number of reads aligning in each type of region. Interestingly, we found that in datasets derived from the polyclonal H3K27ac antibody a higher number of reads fell into enhancer site regions relative to transcription start site regions when compared to the datasets derived from the monoclonal H3K27ac antibody (**Figure 5A**).

**Figure 5.**
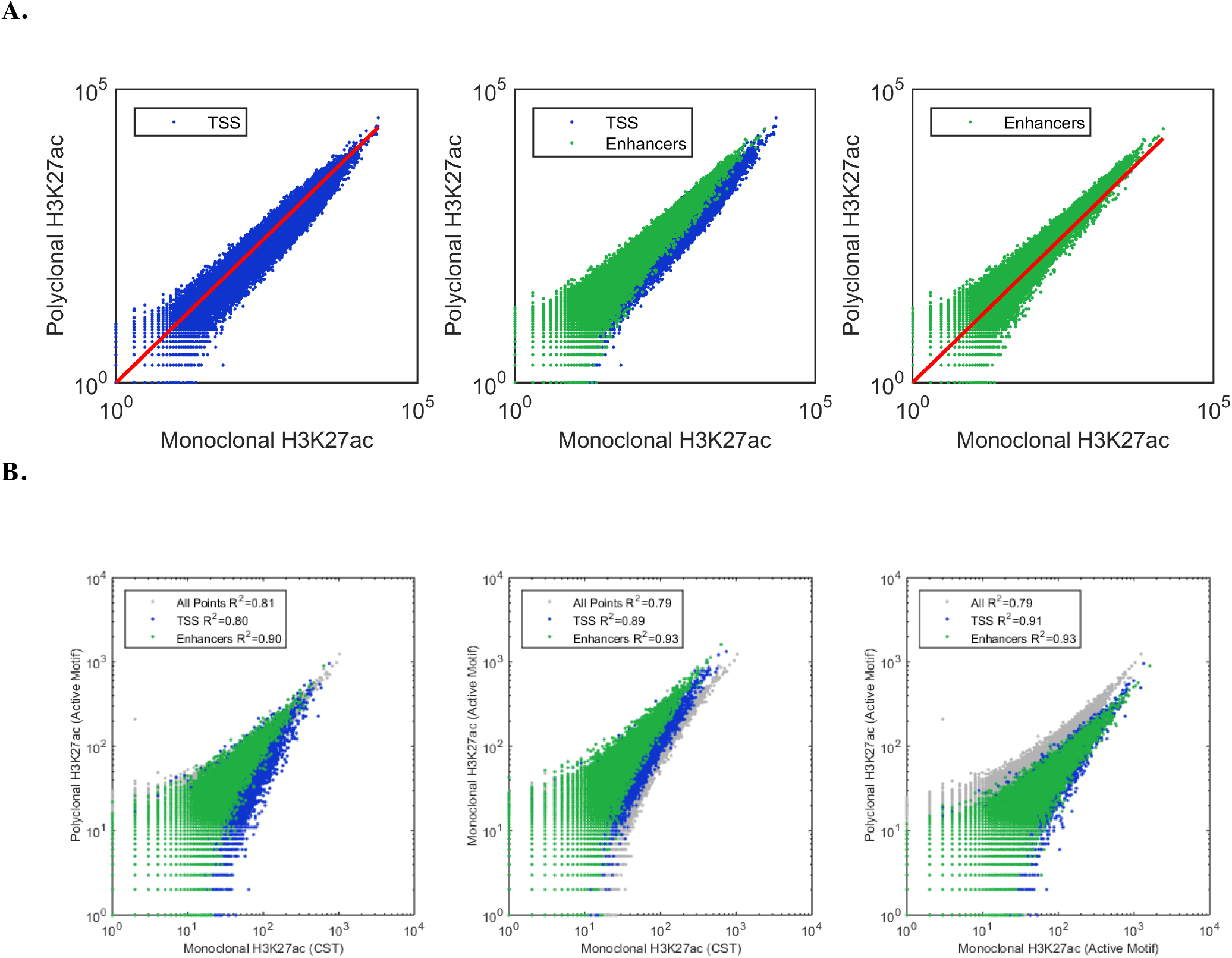
Variability in H3K27ac patterns is dependent on the immunogen. **A.** Scatter plots where each point represents the count of reads aligning to a non-overlapping, variably-sized region as annotated in the chromatin regions determined by ENCODE mapping of the genome. Values are summed for the replicates of monoclonal and polyclonal H3K27ac antibodies. The red line (on the left and right plots) represents slope=1. **B.** H3K27ac antibodies in HeLa cells. R^2^ is indicated for all points, TSS and enhancer regions.

One possible explanation for this observation is that the polyclonal reagent, as it is a mix of individual antibody molecules, contains antibodies to multiple epitopes, one of which is enhancer-specific and increases the antibody’s binding in this region. To examine this possibility, we performed ChIP-seq comparing three H3K27ac antibodies: the monoclonal and polyclonal mentioned above, which were produced by Cell Signaling Technology (CST) and Active Motif, respectively, and a second monoclonal antibody obtained from Active Motif. We repeated this ChIP-seq experiment with the CST monoclonal and polyclonal antibody using HeLa cells and obtained the same pattern (**Figure 5B**). However, when we compared the Active Motif polyclonal antibody to the Active Motif monoclonal antibody the effect was not present. Instead, the ChIP-seq results from the monoclonal Active Motif antibody more closely resembled the polyclonal data (**Figure 5B**).

We were not able to obtain the sequences of the polypeptide immunogens that were used to raise these antibodies as the vendors consider these proprietary. However, the Active Motif antibodies were raised by two different immunogens having an overlapping amino acid sequence (disclosed by Active Motif’s Technical Support to assist with understanding of the data generated for this project). These immunogens likely differed from the one used by Cell Signaling Technology.

### Replication of monoclonal antibodies across lots

To confirm the reproducibility of the monoclonal antibodies between lots, we compared the performance of two different lots of the monoclonal antibody targeting H3K4me3 in K562, GM12878, and mouse ES cells. For each of these lots, we generated two technical replicates and data were normalized by insert size. We quantified the genome-wide performance of the antibody by dividing the genomes into 2000bp bins and counting the reads that aligned to each bin. We found that the correlations between replicates from the same lot were indistinguishable from the correlations across lots (**Figure 6**).

**Figure 6.**
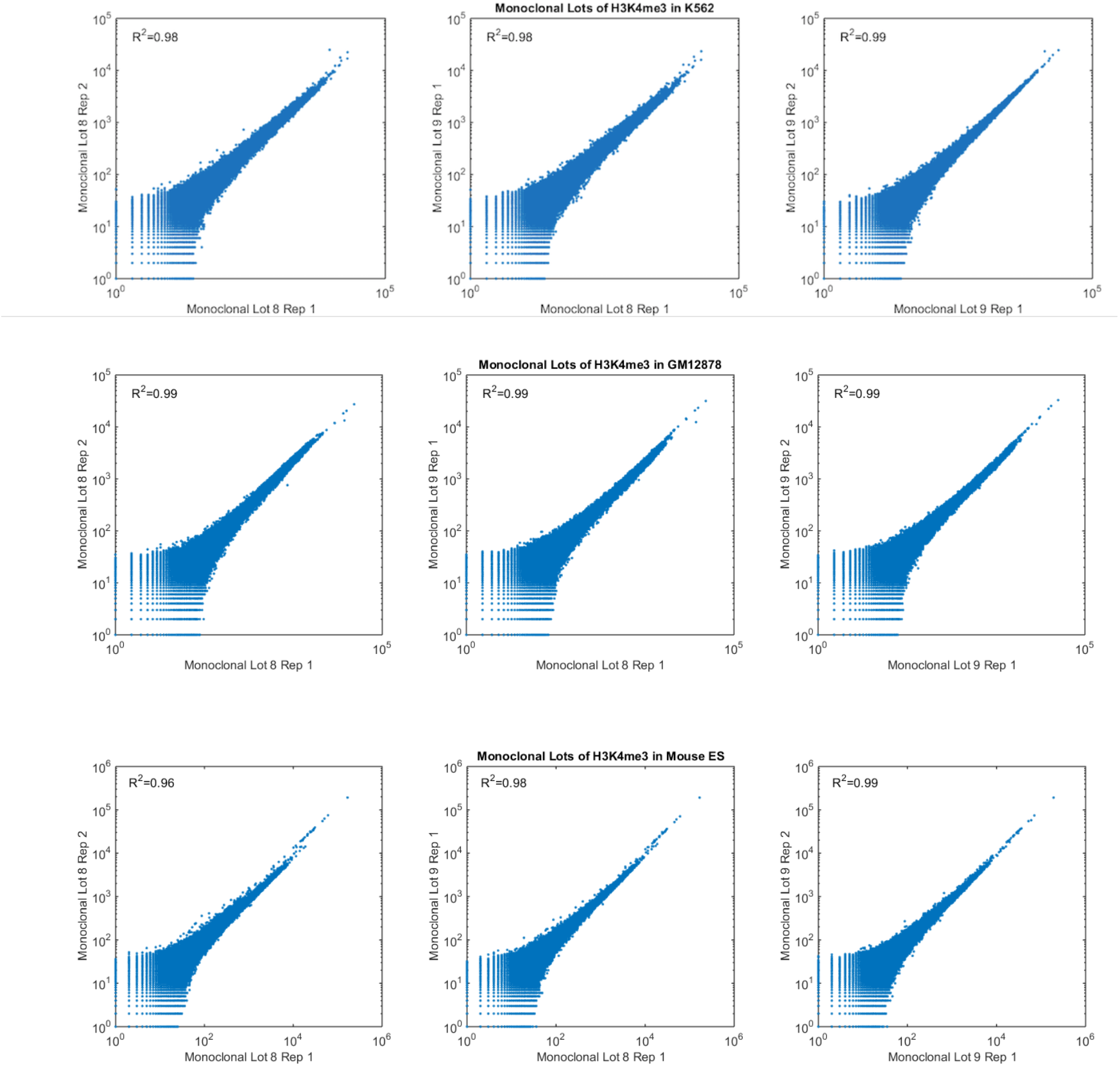
Correlation between two monoclonal lots across the genome. Scatter plots (Loglog) presenting counts of reads per bin in non-overlapping 2000 bp windows tiled throughout the genome comparing either technical or lot replicates from ChIP-seq done with H3K4me3 monoclonal antibody in K562, GM12878 and mES.

### Performance of Monoclonal Antibodies in Other Cell Types

To investigate the performance of the monoclonal antibodies in other cell types, we carried out ChIP-seq in two additional cell lines an EBV transformed human lymphoblastoid cell line (GM12878; **Methods**) and mouse embryonic stem cells. The data demonstrate the performance of these monoclonal antibodies both in a second species and in primary cells that have been shown to have an ‘open’ epigenomic organization.

Next, we compared the performance of the monoclonal and polyclonal antibodies in ChIP-seq using publicly available datasets from the ENCODE consortium from both human [1] and mouse [27]. We aligned our data and the ENCODE datasets to the human (hg19) and mouse (mm9) reference genomes (**Methods**). Initial inspection of the data in a genome browser demonstrated that in these cells as well, there is a high degree of similarity in read coverage between monoclonal and polyclonal antibodies (**Supplemental Figure 2**), even though the polyclonal ChIP-seq datasets were generated by other groups, using different biological samples (GM12878), or even distinct mouse ES cell lines (we used the V6.5 cell line (**Methods**), while the public data was derived from ES-Bruce4 or ES-E14 cell lines [27]).

Next, we calculated the SPOT scores for each of the datasets relative to the peaks called by the ENCODE or Mouse ENCODE consortium. As each experiment was performed with at least two replicates, we were able to perform a t-test to test for statistical differences between our data and the ENCODE data. In the GM12878, only the antibodies to H3K4me3 differed in quality. The data from the monoclonal antibodies had substantially higher SPOT scores than either of the ENCODE datasets, indicating better performance. Among the mouse datasets, the monoclonal antibody for H3K9me3 (p<0.01) and H3K4me1 (p<0.05) performed worse than the polyclonal antibody. All other antibodies performed similarly. (**Table 5**)

**Table 5:**
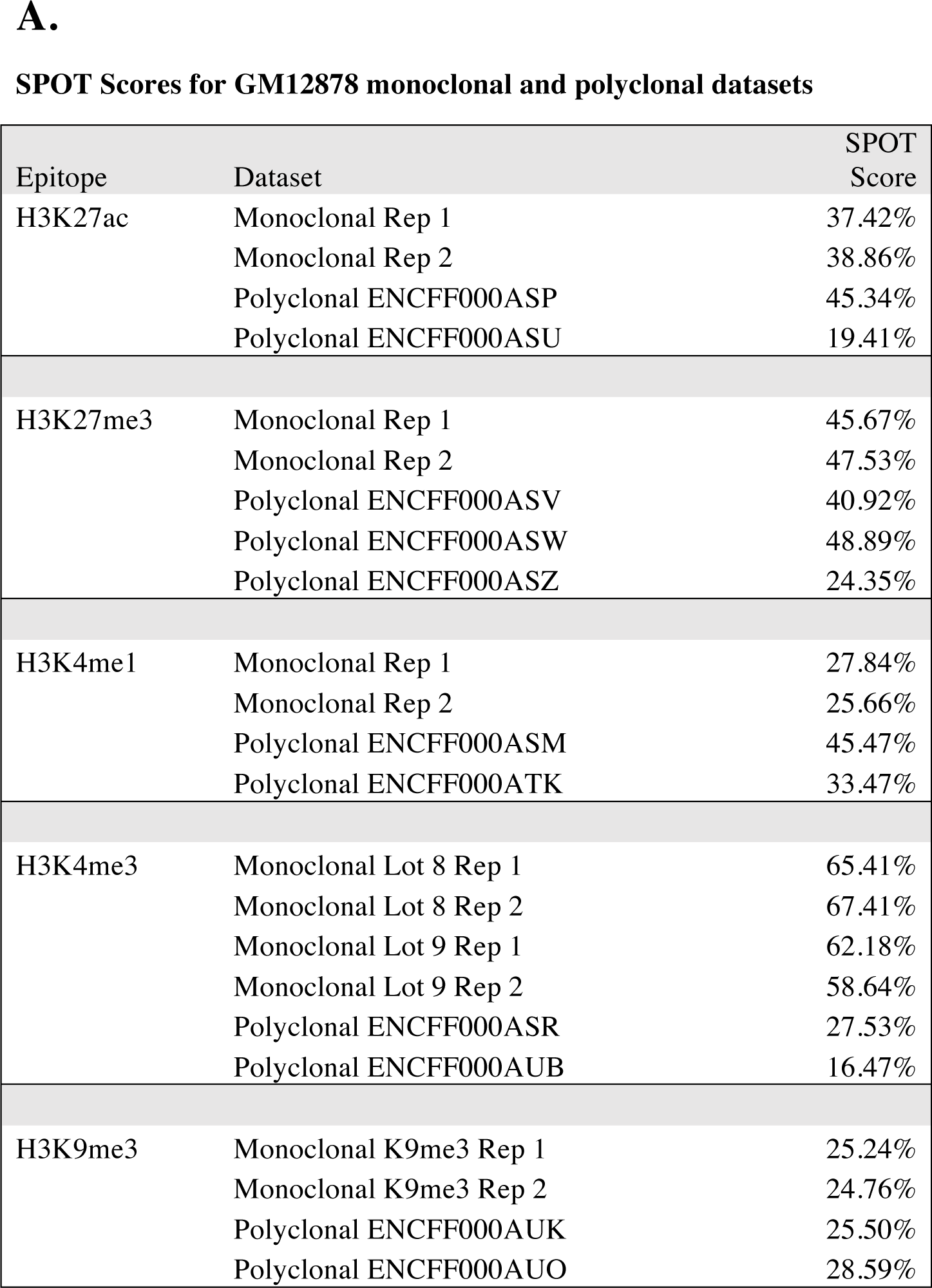
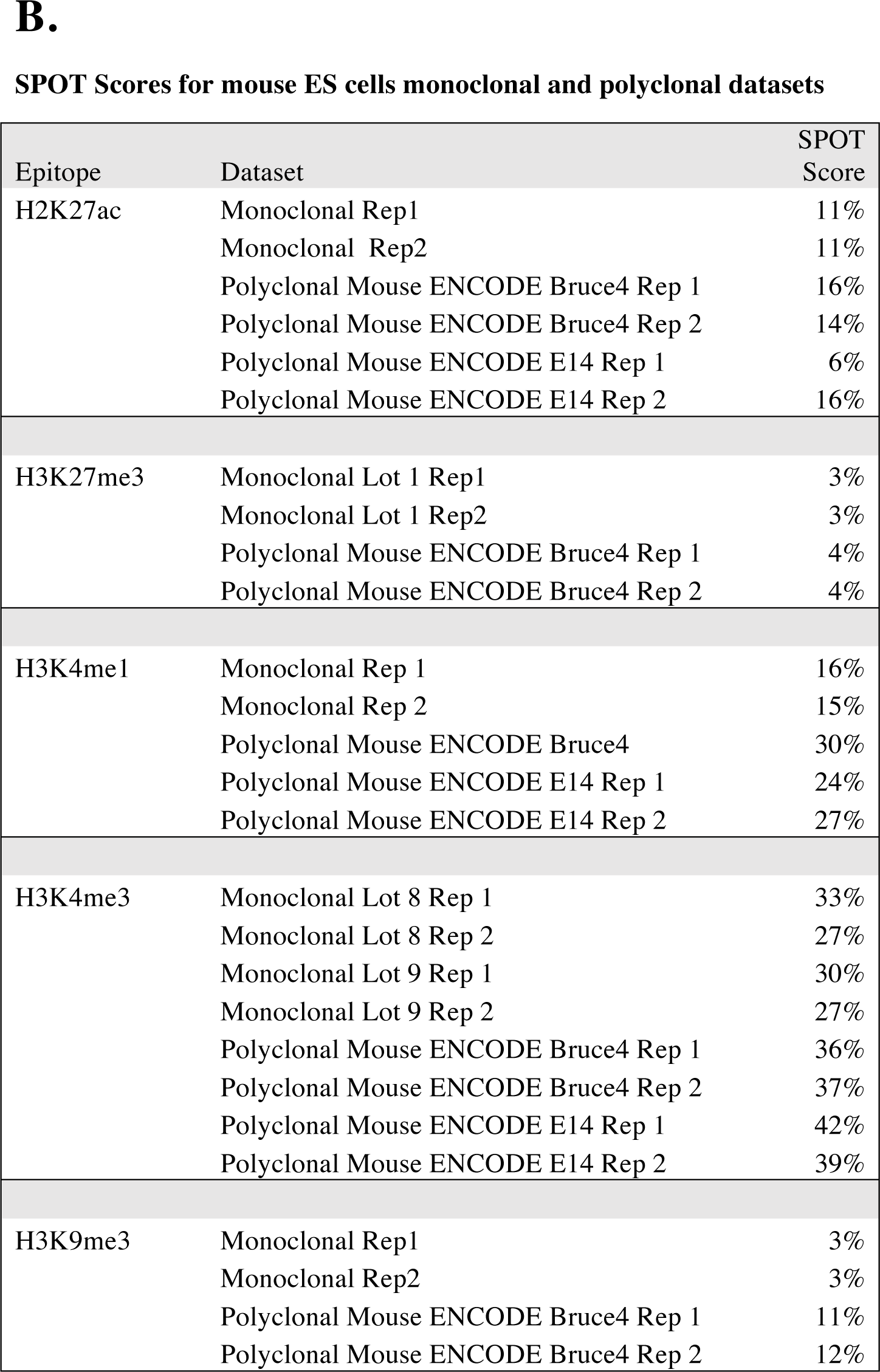
SPOT score for ChIP-seq datasets from GM12878 (**A**) and mouse ES cells (**B**). SPOT score is calculated as the percent of reads that overlap with peaks. For each antibody, peak calls generated by the Mouse ENCODE for the Bruce4 ES cell line (polyclonal antibody) were used to define the peak coordinates.

| Epitope | Dataset | SPOT Score |
| --- | --- | --- |
| H3K27ac | Monoclonal Rep 1 | 37.42% |
|  | Monoclonal Rep 2 | 38.86% |
|  | Polyclonal ENCFF000ASP | 45.34% |
|  | Polyclonal ENCFF000ASU | 19.41% |
| H3K27me3 | Monoclonal Rep 1 | 45.67% |
|  | Monoclonal Rep 2 | 47.53% |
|  | Polyclonal ENCFF000ASV | 40.92% |
|  | Polyclonal ENCFF000ASW | 48.89% |
|  | Polyclonal ENCFF000ASZ | 24.35% |
| H3K4me1 | Monoclonal Rep 1 | 27.84% |
|  | Monoclonal Rep 2 | 25.66% |
|  | Polyclonal ENCFF000ASM | 45.47% |
|  | Polyclonal ENCFF000ATK | 33.47% |
| H3K4me3 | Monoclonal Lot 8 Rep 1 | 65.41% |
|  | Monoclonal Lot 8 Rep 2 | 67.41% |
|  | Monoclonal Lot 9 Rep 1 | 62.18% |
|  | Monoclonal Lot 9 Rep 2 | 58.64% |
|  | Polyclonal ENCFF000ASR | 27.53% |
|  | Polyclonal ENCFF000AUB | 16.47% |
| H3K9me3 | Monoclonal K9me3 Rep 1 | 25.24% |
|  | Monoclonal K9me3 Rep 2 | 24.76% |
|  | Polyclonal ENCFF000AUK | 25.50% |
|  | Polyclonal ENCFF000AUO | 28.59% |
| H2K27ac | Monoclonal Rep1 | 11% |
|  | Monoclonal Rep2 | 11% |
|  | Polyclonal Mouse ENCODE Bruce4 Rep 1 | 16% |
|  | Polyclonal Mouse ENCODE Bruce4 Rep 2 | 14% |
|  | Polyclonal Mouse ENCODE E14 Rep 1 | 6% |
|  | Polyclonal Mouse ENCODE E14 Rep 2 | 16% |
| H3K27me3 | Monoclonal Lot 1 Rep1 | 3% |
|  | Monoclonal Lot 1 Rep2 | 3% |
|  | Polyclonal Mouse ENCODE Bruce4 Rep 1 | 4% |
|  | Polyclonal Mouse ENCODE Bruce4 Rep 2 | 4% |
| H3K4me1 | Monoclonal Rep 1 | 16% |
|  | Monoclonal Rep 2 | 15% |
|  | Polyclonal Mouse ENCODE Bruce4 | 30% |
|  | Polyclonal Mouse ENCODE E14 Rep 1 | 24% |
|  | Polyclonal Mouse ENCODE E14 Rep 2 | 27% |
| H3K4me3 | Monoclonal Lot 8 Rep 1 | 33% |
|  | Monoclonal Lot 8 Rep 2 | 27% |
|  | Monoclonal Lot 9 Rep 1 | 30% |
|  | Monoclonal Lot 9 Rep 2 | 27% |
|  | Polyclonal Mouse ENCODE Bruce4 Rep 1 | 36% |
|  | Polyclonal Mouse ENCODE Bruce4 Rep 2 | 37% |
|  | Polyclonal Mouse ENCODE E14 Rep 1 | 42% |
|  | Polyclonal Mouse ENCODE E14 Rep 2 | 39% |
| H3K9me3 | Monoclonal Rep1 | 3% |
|  | Monoclonal Rep2 | 3% |
|  | Polyclonal Mouse ENCODE Bruce4 Rep 1 | 11% |
|  | Polyclonal Mouse ENCODE Bruce4 Rep 2 | 12% |

### Experimental quality control

To ensure that our ChIP-seq results were representative of the quality of the antibody rather than differences in the performance of the libraries or experiments, one replicate of the H3K27me3 polyclonal antibody was removed as it did not pass our quality control and differed substantially from the other three technical replicates (**Supplemental Figure 6**). Specifically, the number of reads falling into regions of transcription start sites was systematically higher in this replicate than in other replicates. A monoclonal replicate of the H3K4me1 and a monoclonal replicate of H3K9me1 failed to yield an adequate number of reads to be used in analysis. These samples were rerun in duplicate and each was replaced with two replicates.

## Discussion

Our goal in designing this study was to improve current ChIP-seq procedures by increasing the reproducibility between experiments within the community, as well as to enhance the usage of reagents that have long-term accessibility. Specifically, we explored whether monoclonal antibodies could properly replace the polyclonal antibodies routinely used in ChIP-seq for detection of histone post-translational modifications.

Our experimental design allowed us to directly compare performance of monoclonal and polyclonal antibodies in ChIP-seq assays. First, we used standardized automation for all laboratory processes. This virtually eliminated variation arising from human handling and ensured that all samples were handled as identically as possible. Second, all sequence data for this study were generated as paired-end reads, since paired-end data provide the only definitive means to assess the lengths of DNA fragments that were sequenced. Accordingly, we were able to leverage the paired-end data to normalize alignments to eliminate fragmentation and size selection biases as confounding factors. We observed a high degree concordance between results from data normalized by insert size and from data normalized for read coverage depth by random downsampling. Thus, differences in fragmentation and size selection did not appear to be confounders in this work.

Our analysis demonstrated that the insert length of paired-end reads correlated with the genomic regions from which the fragments originated, consistent with earlier reports [21]. We therefore strongly recommend optimizing fragmentation and size selection protocols to include the full range of genomic fragment sizes to avoid bias, as well as using paired-end reads for ChIP-seq experiments to account for variation in fragment size among samples and to allow accounting for amplification-based duplicates in the sequencing libraries. In future studies, it would be useful to evaluate whether insert size normalization can provide a cost-effective alternative to using WCE controls, particularly in experiments whose primary focus is to measure changes in protein binding under different conditions rather than an exhaustive mapping of binding locations.

Among the five antibodies tested, the polyclonal antibodies to H3K4me3 and H3K27me3 appeared to offer slightly higher sensitivity while the monoclonal antibody to H3K27ac appeared to offer higher specificity. However, the differences in H3K27ac are more likely result from the specific immunogen against which the antibody was raised rather than the clonality of the antibody. Because higher sensitivity was not seen in the other polyclonal antibodies, our results demonstrate that the use of monoclonal antibodies for ChIP-seq did not present any systematic disadvantage relative to polyclonal antibodies, and have the clear advantage of superior reproducibility. This conclusion is supported by high correlation in both genome-wide and region-specific read counts between monoclonal and polyclonal antibodies, as well as the high degree of overlap in peak locations in multiple cell types and two distinct species.

Overall, our data are consistent with a model suggested by Peach and colleagues [28] in which some antibodies are better described as indicators of canonical regions of the genome rather than as markers of specific modifications. For instance, in our comparison of H3K27ac antibodies, the monoclonal and polyclonal antibodies displayed significant differences in their relative ratios of reads localized to putative enhancers versus transcription start sites. If we assume that the targeted acetylated H3K27 is the same molecule in each region, then the ability of the antibodies to identify H3K27ac was affected not just by the presence of the target but also by its local environment. This finding is expected as characteristics of the environment, such as neighboring post-translational modifications, have been demonstrated to detectably affect epitope recognition [4]. The binding pattern of a single antibody should thus be thought of as a collection of component parts that describe more than just the binary presence or absence of a modification. This inherent complexity is further complicated by the fact that researchers often do not know the precise nature of the immunogen that was used to raise a specific antibody because the antibody’s producer holds this information as proprietary.

Thus, ChIP-seq datasets targeting the same epitope but using different antibodies cannot be considered directly comparable without substantial experimental validation. Standardizing on monoclonal antibodies would not only eliminate the batch-to-batch variability that is expected in polyclonal antibodies but would also increase the value of ChIP-seq datasets by allowing for more reliable reuse of existing datasets. Further, it would simplify the interpretation of ChIP-seq data by removing the added complexity that is introduced by using a polyclonal antibody that targets an unknown number of epitopes on the antigen.

The relative portion of reads aligned to different canonical regions of the genome was also affected by experimental variability. By examining the relative proportion of reads mapping to canonical regions of the genome, we were able to easily identify an outlier replicate in our H3K27me3 data that would have passed less rigorous quality standards. This finding demonstrates not only that replicates are imperative in any ChIP-seq experiment, but also that performing this simple analysis can provide valuable information for quality control.

## Conclusions

Use of monoclonal antibodies for ChIP-seq experiments to identify histone post-translational modifications provides a key improvement over polyclonal ones.

## Methods

### Chromatin Immunoprecipitation (ChIP)

ChIP comprises the basic steps of crosslinking DNA to protein, shearing DNA, and enriching of the protein of interest, along with DNA to which is it crosslinked, by immunoprecipitation. Washes and mixes were conducted using the Bravo liquid handling platform (Agilent model 16050-102, “Bravo”). For the compositions of the buffers used, see [29]; for the specific protocol for the Bravo, see [19].

**Table 1** describes the antibodies used in this study. The polyclonal antibodies, including these specific lots, were previously assessed for accuracy by the ENCODE consortium.

*Crosslinking and DNA Shearing:* K562 mylogenous leukemia cells (ATCC CCL-243), GM12878 lymphoblastoid cells (Coriell; Cellosaurus GM12878 (CVCL_7526)) and mouse ES cell line V6.5 (Cellosaurus v6.5 (CVCL_C865)) were cross-linked with formaldehyde as previously described [29]. Fixed cell pellets (20 million cells each) were resuspended in lysis buffer and ChIP dilution buffer and incubated on ice to lyse the cells. Samples were then split across a 96-well plate (approximately 1–2 million cells per well). DNA shearing was conducted using a Covaris sonifier (model E220) at 4°C for 6 cycles of 1 minute, with these parameters DF-10%, PIP-175W, CPB-200. After sonication, the cell lysates were diluted 1:10 with ChIP dilution buffer. Roughly 50 µL of the cell lysate was set aside for use as the whole cell extract (WCE) control.

*Bead Preparation:* Immunoprecipitation was performed using magnetic beads coupled to antibodies by Protein A or Protein G linkers. The beads were prepared as follows: equal quantities of Protein A and Protein G Dynabeads (Invitrogen, 100-02D and 100-07D, respectively) were mixed, separated into 50 µL aliquots in a well plate, and washed twice with blocking buffer. The beads and antibodies (5 µL of polyclonal or 1 µL monoclonal antibody per ChIP reaction), mixed and suspended in blocking buffer, were incubated in a cold room (4°C) on a rotator for at least two hours to allow conjugation.

*Immunoprecipitation of Target Protein and DNA Purification:* Washed bead-antibody conjugates were added to the chromatin lysate from approximately 1–2 million cells and incubated overnight. At this point, the WCE was added to the sample plate. Samples were washed six times with RIPA buffer, twice with RIPA buffer supplemented with 500 mM NaCl, twice with LiCl buffer, twice with TE, and then eluted in ChIP elution buffer to unlink and purify the DNA.

### Library Construction

The library construction phase of ChIP-seq comprises DNA end-repair, A-base addition, adaptor ligation, and enrichment. Solid-Phase Reversible Immobilization (SPRI) cleanup was performed on the reverse-crosslinked DNA before library construction and after each of its four steps to remove proteins and other molecules.

*SPRI Cleanup Protocol:* SPRI cleanup steps were conducted using the Bravo, following protocols described by [19]. All enzymes used in library construction were obtained from New England Biolabs. The initial and final SPRI cleanups for the reverse-cross-linked DNA were performed as follows: SPRI beads (Agencourt AMPure XP) were added to the unlinked DNA samples. The beads were washed on a 96-well bar magnet (ThermoFisher, catalog number: 12027) with 70% ethanol and air-dried. The DNA was eluted in 10 mM Tris-HCl buffer. Intermediate SPRI cleanups in the library construction process were conducted in the same manner. The SPRI beads in the reaction were reused to capture the DNA via addition of a 20% PEG solution.

*End-Repair and A-base Addition:* DNA end-repair was performed by adding T4 PNK enzyme and T4 polymerase to each well, followed by incubation at 12°C for 15 minutes and at 25°C for another 15 minutes. Following SPRI clean up, A-base addition was performed by adding Klenow 3’ → 5’ exonuclease and incubation at 37°C for 30 minutes.

*Adapter Ligation:* Adapter ligation was performed by adding DNA ligase and PE Indexed oligonucleotide adapters to samples followed by incubation at 25°C for 15 minutes. After the subsequent SPRI cleanup, eluted DNA was separated from the SPRI beads using a 96-well bar magnet for PCR enrichment.

*Enrichment:* DNA samples were PCR amplified at 95°C for 2 minutes; 16 cycles of: 95°C for 30 seconds, 55°C for 30 seconds, 72°C for 60 seconds; and 72°C for 10 minutes.

### Data Collection and Analysis

DNA fragments were processed by 2x25 base, paired-end or 2x37 base, paired-end sequencing (Illumina HiSeq 2500 or NextSeq 500, respectively).

To assess reproducibility, we designed an analysis pipeline consisting of the following steps: alignment, normalization, pairwise correlation and clustering, peak calling, and analysis. Reads were aligned by the Broad Genomics Platform with BWA (v5.9) using default parameters [30].

To allow for meaningful comparisons between different samples, duplicate reads were removed from the alignment data (BAM file) using the Picard tools software package. Downsampling was performed using C++ scripts built using the BamTools API [31]. Scripts are available on GitHub (https://github.com/mbusby/).

Downsampling normalization by insert size was performed as follows: We first counted how many read pairs are present for each insert size for each of a set of aligned files. We then selected the lowest read count for each insert size from among the set of alignments. For example, if four alignments for a given antibody have one, two, three, and four million reads with an insert size of 100, all four alignments would be randomly sampled so that the four normalized alignments each have about one million reads with an insert size of 100. This was performed for each insert size present in all of the alignments in the group to yield final bam files with about the same numbers of reads and insert size distributions. This approach therefore allows for identical insert size distributions while maximizing the number of reads included in the output files. All samples for each histone modification were sampled as a group. The K562 WCE control and the HeLa samples were not downsampled. The merged datasets used in peak calling were created by merging the technical replicates downsampled by insert size. To create balanced datasets, in cases where one antibody had more replicates than its counterpart an equal number of replicates were used for the monoclonal and polyclonal dataset. Replicates were chosen based on the order of their replicate number.

Peaks were called using HOMER (v.4.7) [24] with the WCE used as a control under the default settings for paired-end reads using “histone” as the peak type.

We used the BEDtools coverage tool, version 2.25 [32] to count the number of reads mapping to genomic regions and the intersect tool to count the genomic reads that overlapped between antibody types. The combined Segway and ChromHMM annotations were downloaded from [33]. Further analyses were performed in Matlab. Scripts are available on GitHub (https://github.com/mbusby/).

## Declarations

### Abbreviations

ChIP-seq: Chromatin immunoprecipitation followed by sequencing
CST: Cell Signaling Technology
WCE: whole cell extract

## Ethical Approval and Consent to participate

Not applicable

## Consent for publication

Not applicable

## Availability of supporting data

Pending submission to SRA.

## Competing Interests

The authors declare no competing interests.

## Funding

This work was supported by The Broad Institute SPARC (Scientific Projects to Accelerate Research and Collaboration) program.

## Authors’ contributions

M.B. and AG designed the experiments; M.B., C.X., C.L., Y.F. and A.G. analyzed and interpreted the data; C.X., C.L., E.G., I.Y., A.Gl., C.B.E., E.M.C. and S.B.R. carried out experiments and provided supporting data; M.B., C.N. and A.G. wrote the manuscript with help from C.X. and E.G.; C.B.E., C.N. and A.G. acquired funding and resources; A.G. supervised the study. All authors have read and approved the final manuscript.

## Acknowledgements

We would like to thank Chip Stewart at the Broad Institute for helpful discussions, and the Broad Institute Genomics Platform for generating all DNA sequence data described here.

---

**Supplemental Figure 1.**
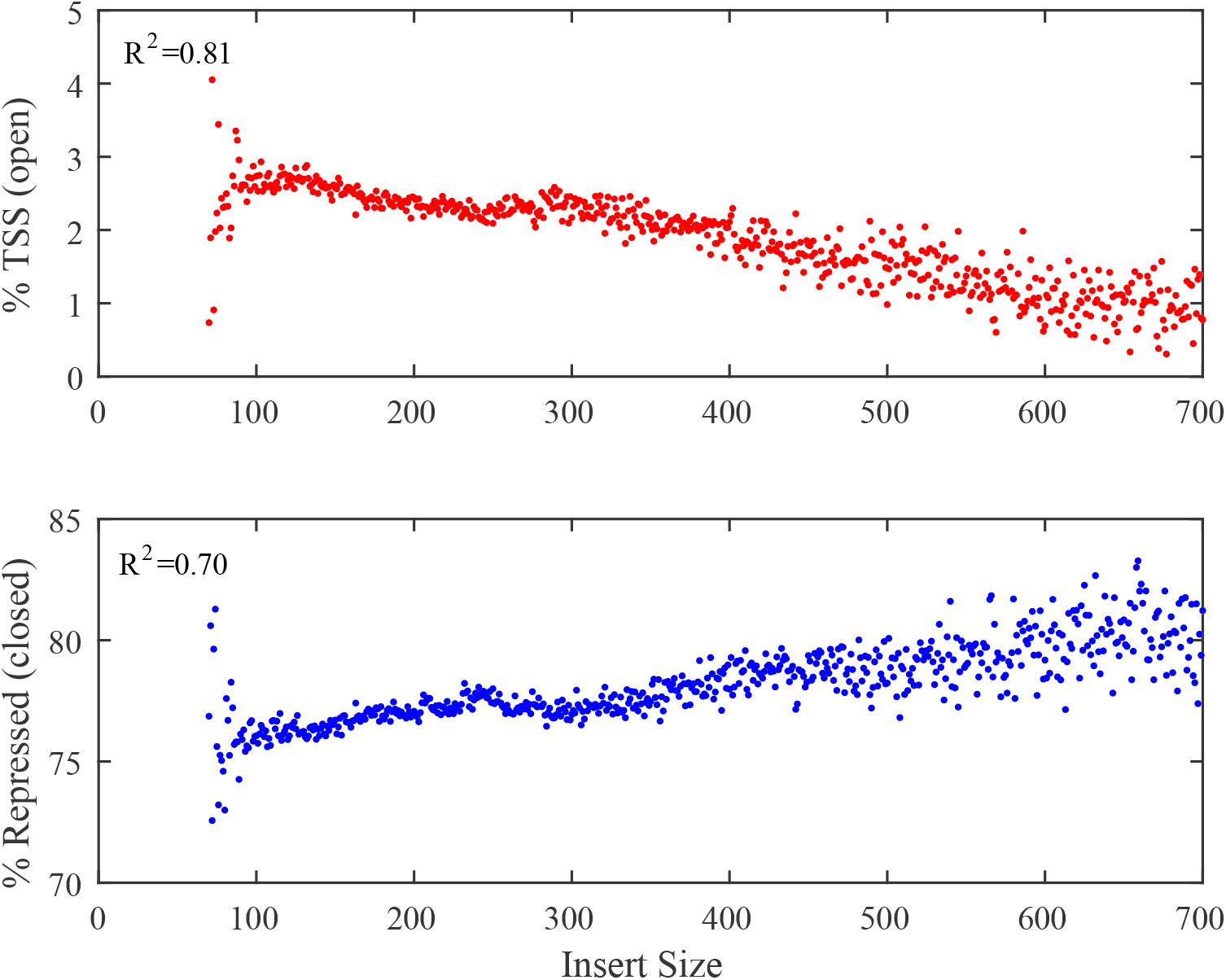
Reads aligning to annotated open and closed chromatin. The location of reads of different insert sizes from our K562 WCE data was compared to the ENCODE annotation of the genome using BEDTools [1]. Reads with smaller insert sizes, indicating that they originate from smaller DNA fragments, mapped to transcription start sites (open chromatin) at a higher rate than those of longer insert sizes. An opposite relationship was oboserved for closed chromatin. Note, however, that the Y axis is truncated to show this effect and the change in the percentage of reads tracking to repressed regions is small. Reads below and above the displayed insert sizes were excluded due to low read coverage and associate noise.

**Supplemental Figure 2.**
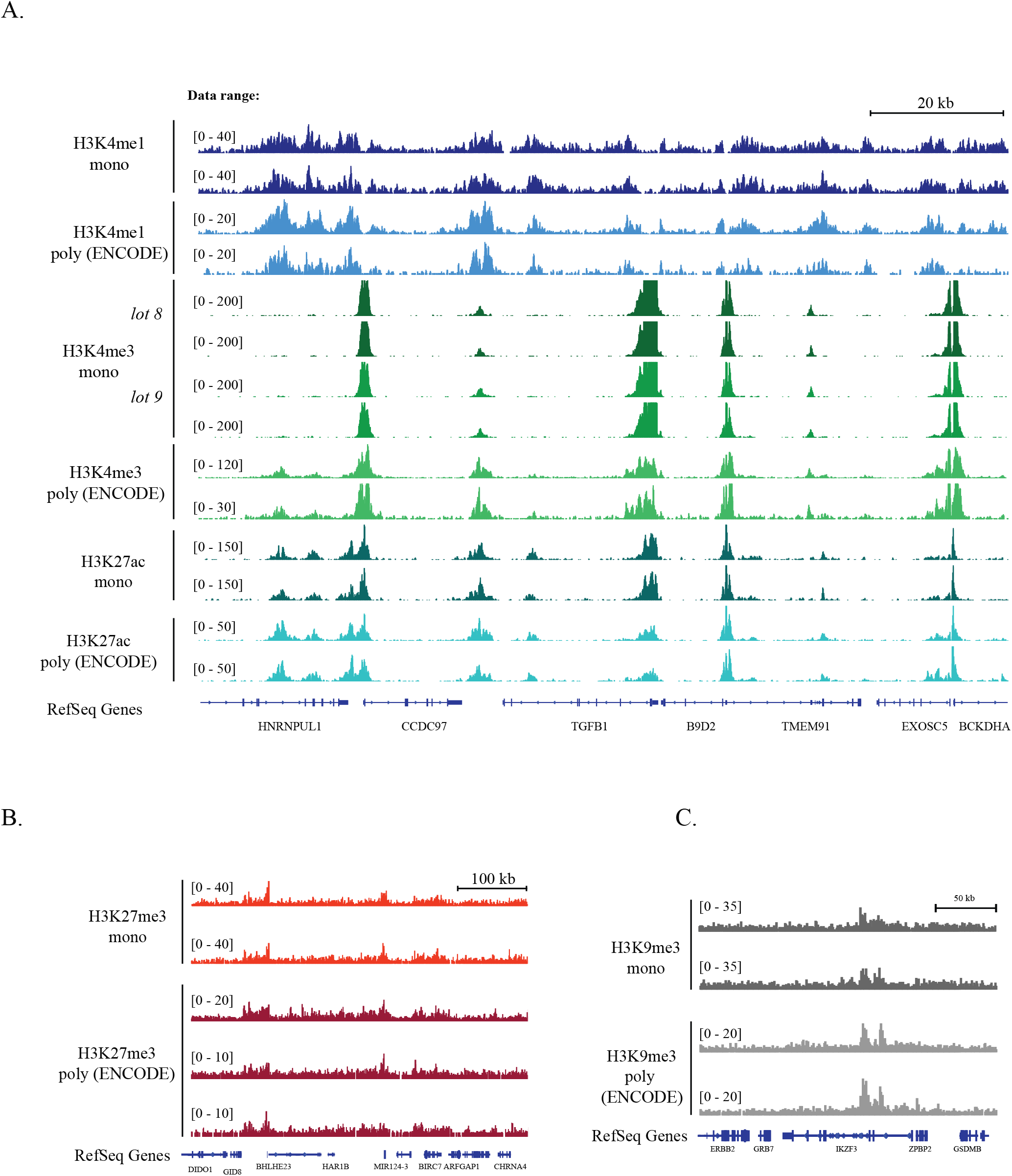
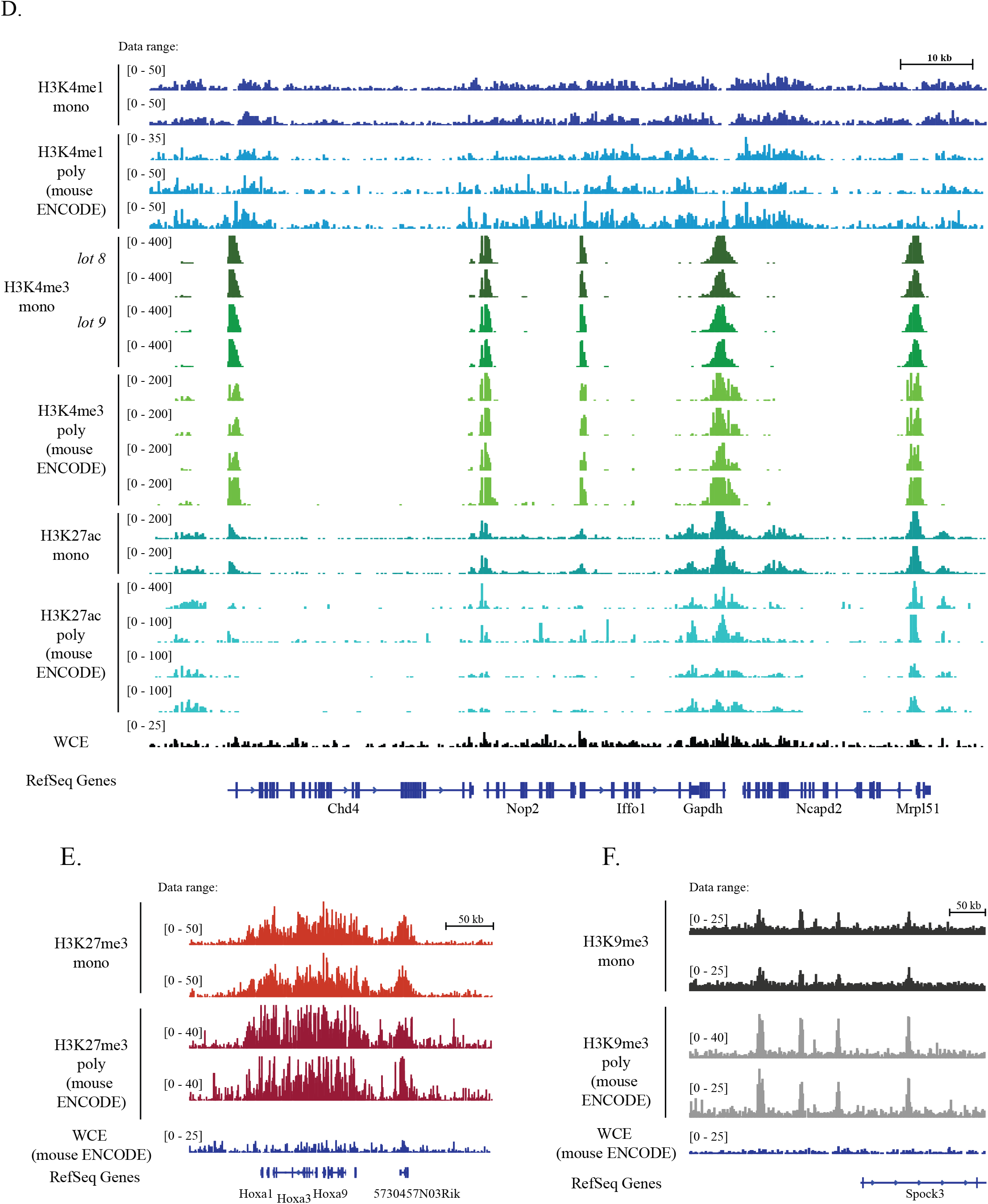
Read coverage across the genome. Images of Tiled Data Files (TDFs) generated by the IGV browser [2, 3] displaying the density tracks of reads aligned across the genome. The tracks show the correspondence in read coverage in monoclonal and polyclonal antibodies (from ENCODE and Mouse ENCODE) over representative genomic loci. **A-C** GM12878; **D-F** mES. **A.** Chromosome 19:41,792,290-41,912,756 (about 120Kb) for H3K4me1, H3K4me3 (two monoclonal lots) and H3K27ac. **B.** Chromosome 20:61,503,810-62,032,504 (about 500Kb) for H3K27me3. **C.** Chromosome (about 250Kb) for H3K9me3. **D.** Chromosome 6:125,035,929-125,153,426 (about 120Kb) for H3K4me1, H3K4me3 (two monoclonal lots) and H3K27ac. **E.** Chromosome 6:51,949,208-52,399,454 (about 450Kb) for H3K27me3. **F.** Chromosome 8:65,216,638-65,604,437 (about 390Kb) for H3K9me3.

**Supplemental Figure 3.**
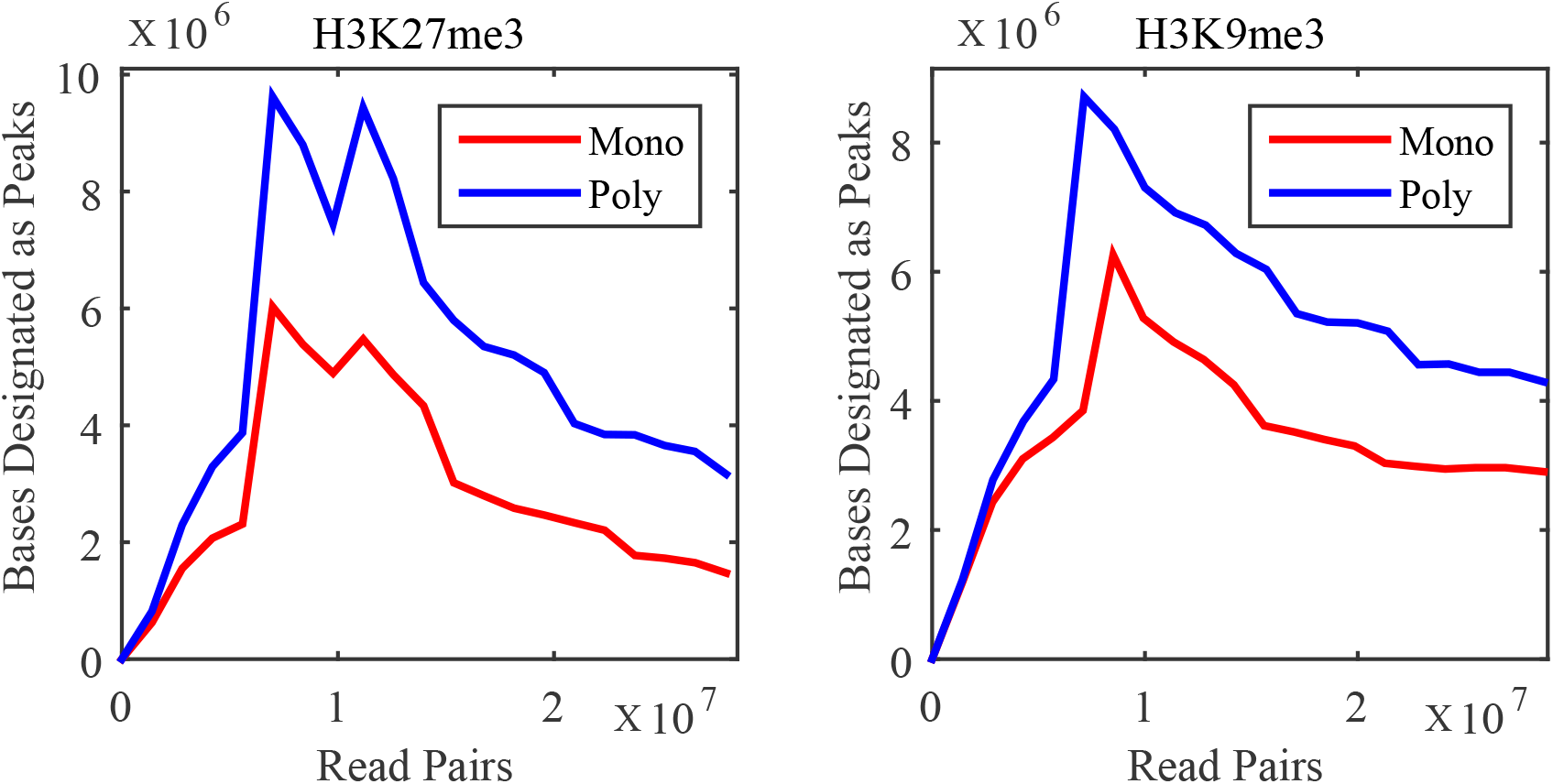
Saturation curve showing the number of bases called as being in peaks as a function of sequencing depth for H3K9me3 and H3K27me3. The final dataset of the merged technical replicates was randomly down sampled to 20 different read depths and peaks were called in each dataset using HOMER. For these datasets, curves do not show the expected pattern and do not appear to reach saturation.

**Supplemental Figure 4.**
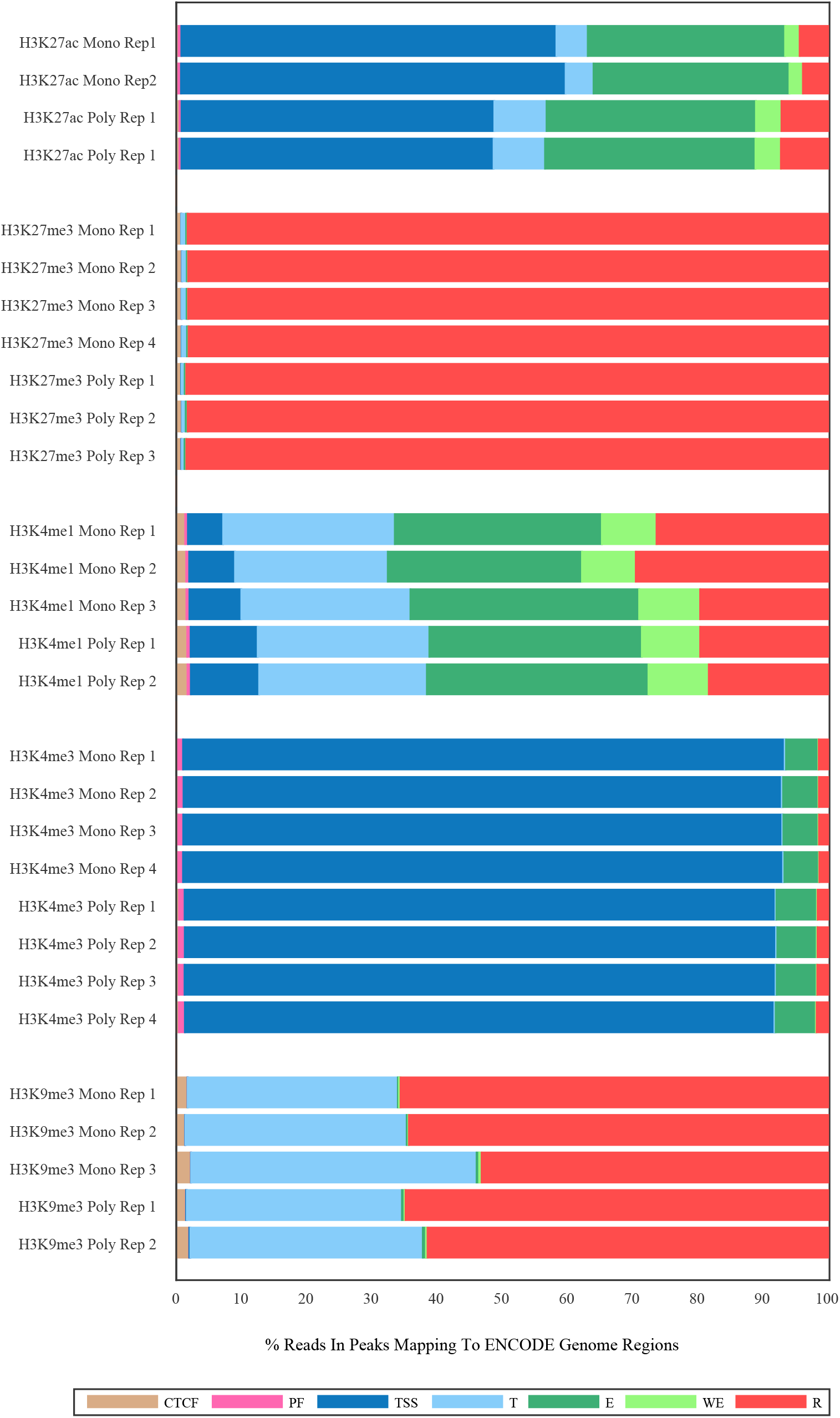
Mapping of peaks and reads to canonical chromatin regions of the genome as defined by the ENCODE mappings. These plots display the percentage of reads that map to each canonical genome region. The canonical genome regions were defined by the combined ENCODE mapping and are abbreviated as follows: CTCF-enriched elements (*CTCF*), promoter flanking regions (*PF*), transcription start sites (*TSS*), transcribed regions (*T*), enhancers (*E*), weak enhancers (*WE*), and repressed regions (*R*). Reads were normalized by random downsampling.

**Supplemental Figure 5.**
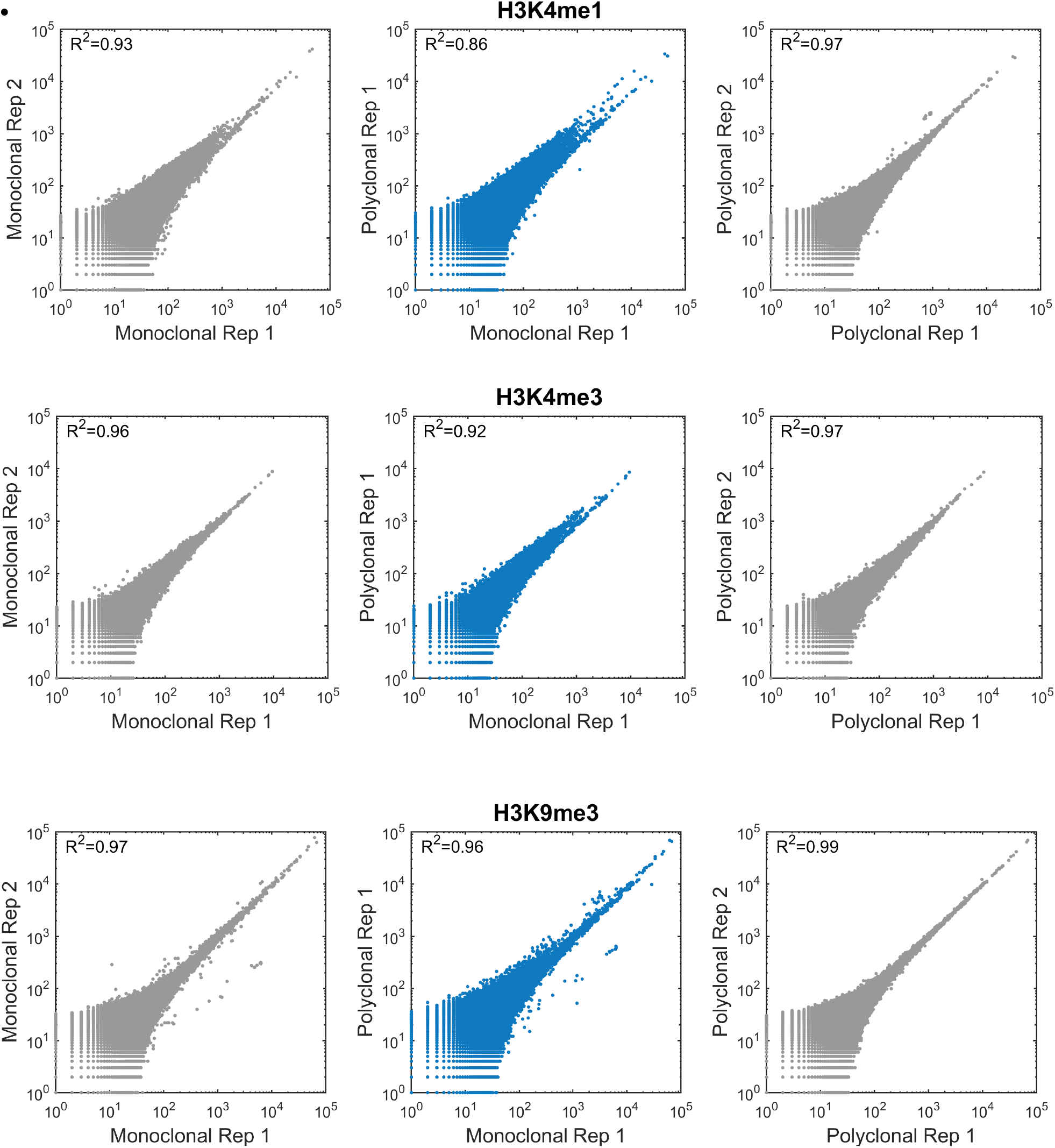
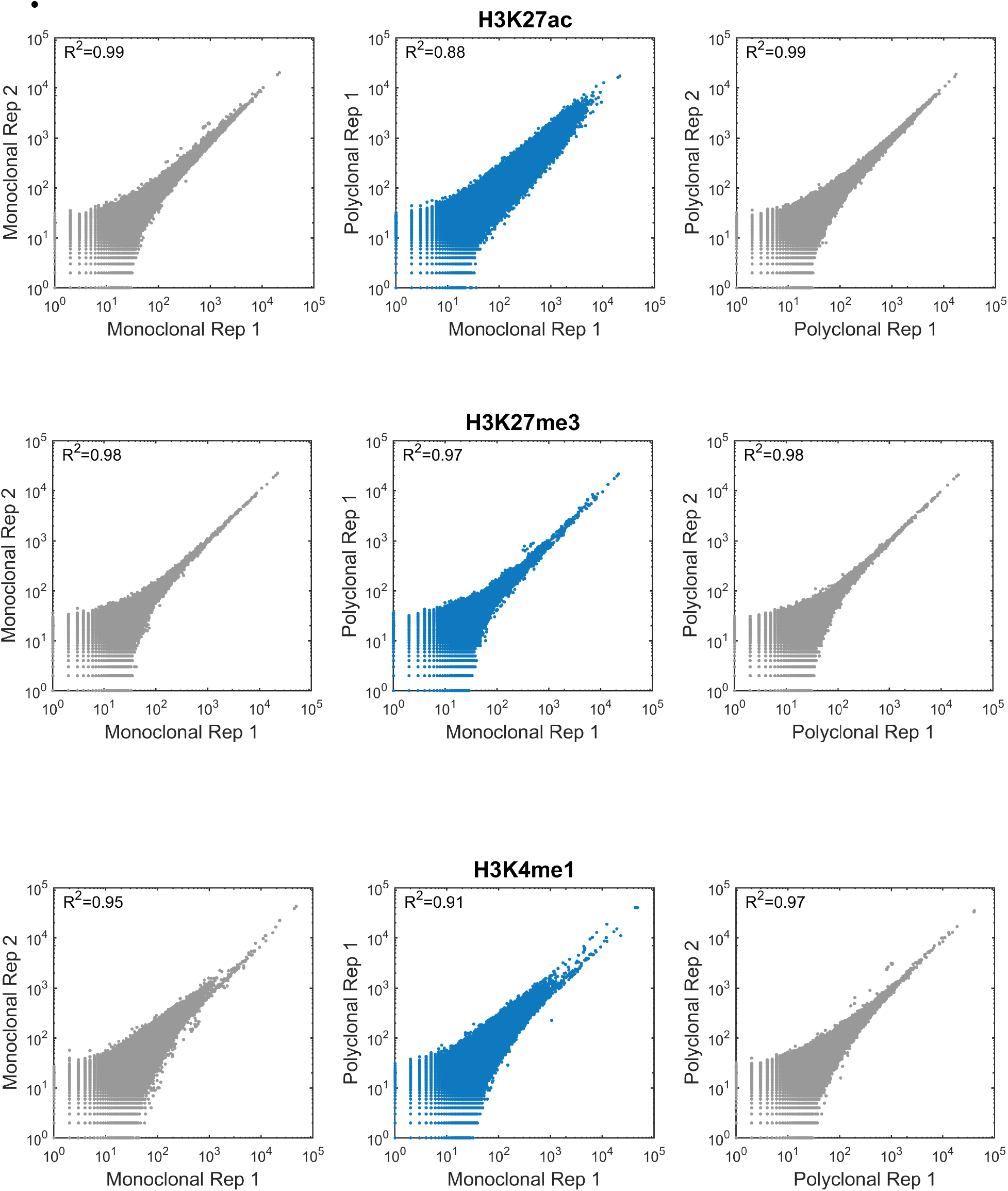
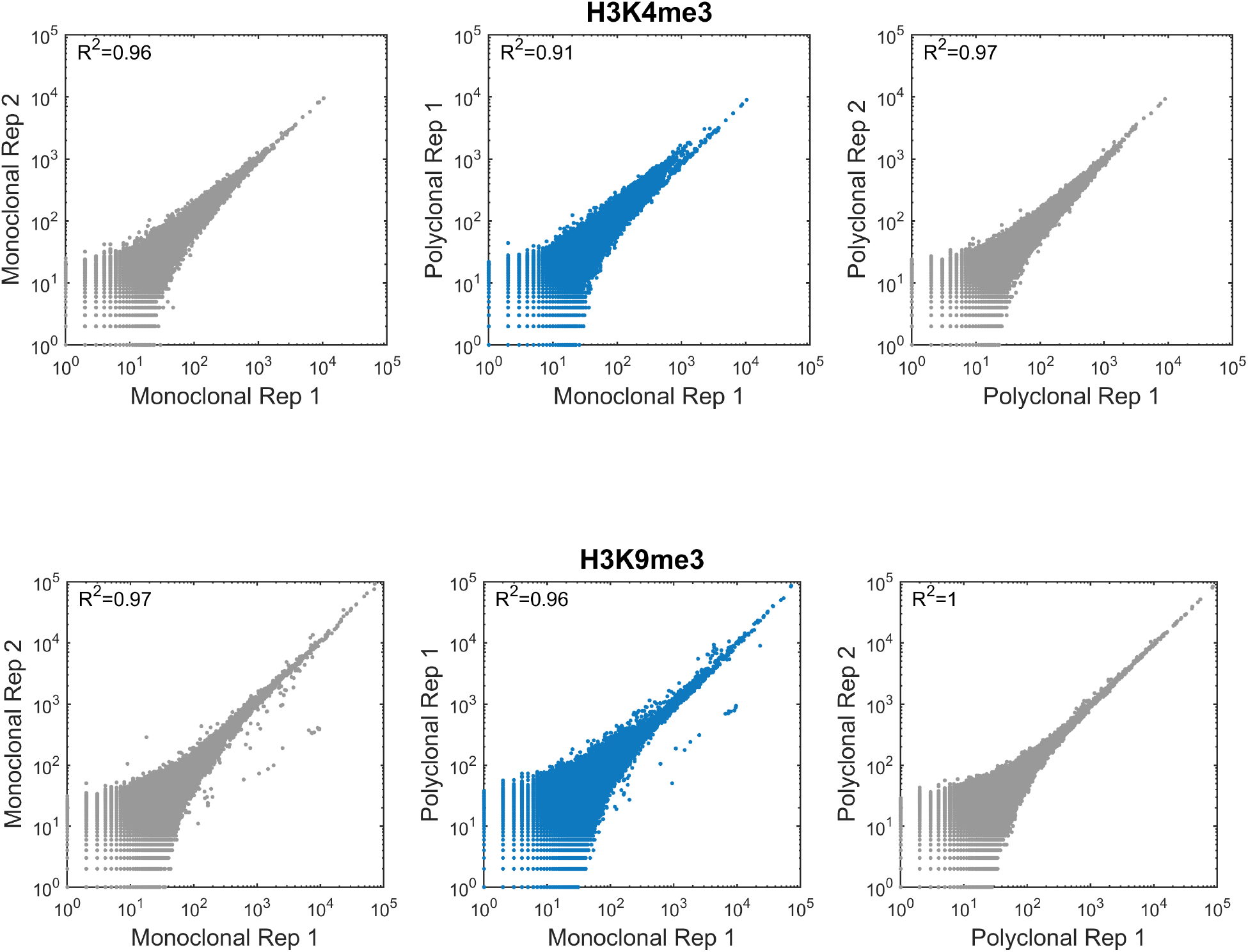
Correlation between monoclonal and polyclonal antibodies across the genome. Scatter plots (Loglog) show counts of reads per bin in non-overlapping 2000 bp bins tiled throughout the genome in replicates of the monoclonal antibody (**left**), the polyclonal antibody (**right**), and polyclonal antibody versus monoclonal antibody (**center**). The reads for each dataset were normalized by **A.** insert size or by **B.** random downsampling.

**Supplemental Figure 6.**
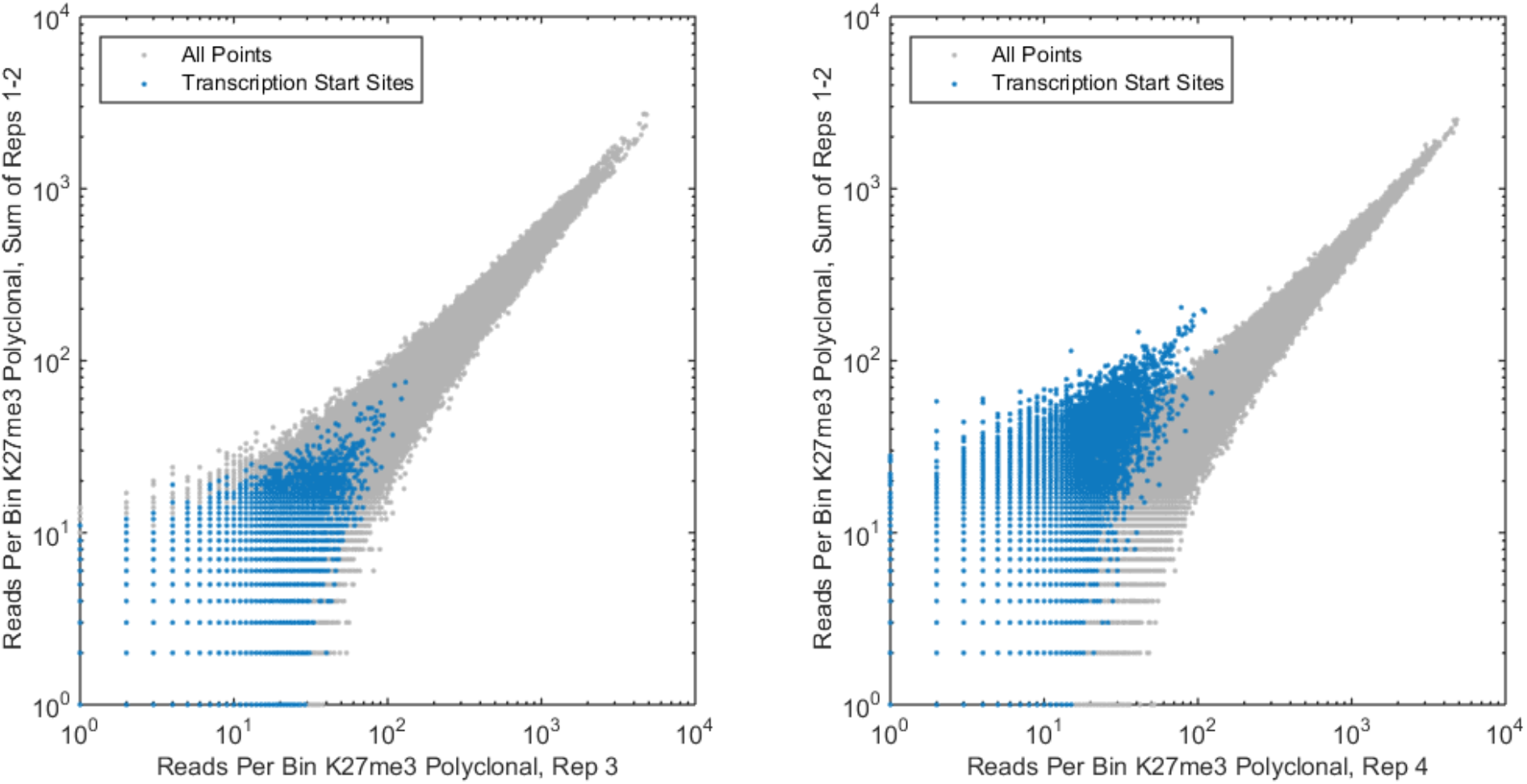
Experimental quality control. While we generated four technical replicates of the H3K27me3 polyclonal antibody, one of the replicates (Replicate number 4) did not pass our quality control. Points on the graph represent the number of reads falling into each variable-sized bin defined by the canonical chromatin regions of the genome as defined by the ENCODE. **A.** A comparison of Replicate 3 to the summed read counts of Replicates 1 and 2. This is in line with our expectations. **B.** The same comparison to Replicates 1 and 2, this time using Replicate 4. Here we see that Replicate 4 has a systematically reduced read count in transcription start sites. No other classes of genomic regions were shifted. We concluded that Replicates 4 did not pass our quality control.

**Supplemental Figure 7.**
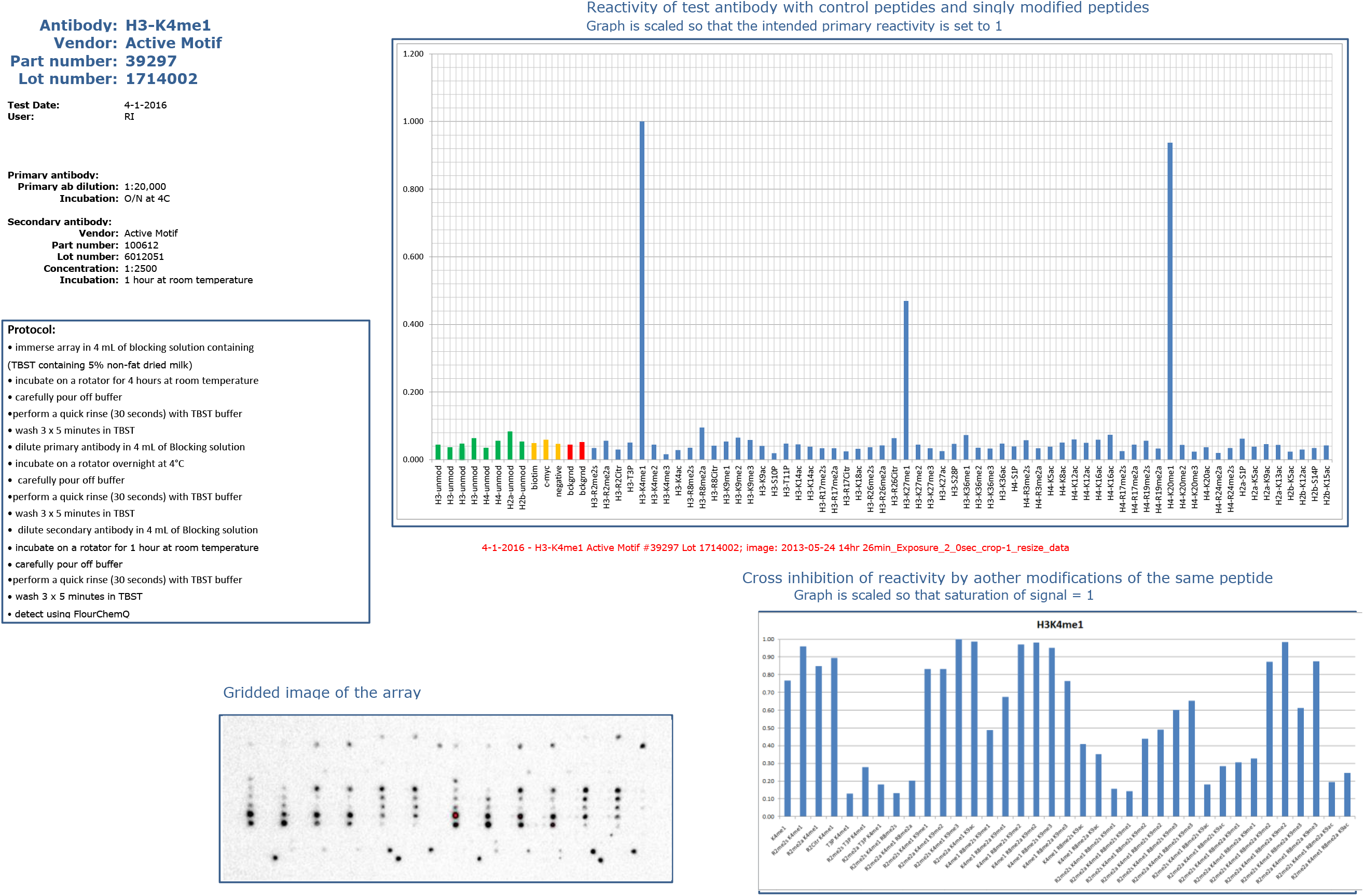
Validation of the polyclonal antibody targeting H3K4me1 by peptide array. The details are presented in the same manner as the publically available validations of the other antibodies we used in the study (**Table 1**), and include the details of the antibody and validation protocol, the row signal of the peptide array (**bottom left**); reactivity with the synthetic peptides on the array (**top right**), and the cross reactivity other modifications of the same peptide.

**Supplemental Table 1.** Datasets summary

| Cell Type | Antibody | Replicate | Reads | Pairs |
| --- | --- | --- | --- | --- |
| GM12878 | H3K4me1 | Monoclonal Rep 1 | 83,100,486 | 41,550,243 |
|  |  | Monoclonal Rep 2 | 99,255,196 | 49,627,598 |
|  | H3K4me3 | Monoclonal Lot 8 Rep 1 | 99,255,196 | 78,227,985 |
|  |  | Monoclonal Lot 8 Rep 2 | 156,455,970 | 42,980,622 |
|  |  | Monoclonal Lot 9 Rep 1 | 85,961,244 | 33,912,720 |
|  |  | Monoclonal Lot 9 Rep 2 | 67,825,440 | 36,033,346 |
|  | H3K9me3 | Monoclonal Rep 1 | 143,527,964 | 71,763,982 |
|  |  | Monoclonal Rep 2 | 66,206,100 | 33,103,050 |
|  | H3K27ac | Monoclonal Rep 1 | 146,244,844 | 73,122,422 |
|  |  | Monoclonal Rep 2 | 156,042,348 | 78,021,174 |
|  | H3K27me3 | Monoclonal Rep 1 | 157,892,610 | 78,946,305 |
|  |  | Monoclonal Rep 2 | 152,454,660 | 76,227,330 |
| HeLa | H3K27ac | Monoclonal Rep 1 | 17,340,796 | 8,670,398 |
|  |  | Monoclonal Rep 2 | 16,389,304 | 8,194,652 |
|  |  | Polyclonal Rep 2 | 13,155,478 | 6,577,739 |
| K562 | H3K4me1 | Monoclonal Rep 1 | 44,707,286 | 22,353,643 |
|  |  | Monoclonal Rep 2 | 19,370,594 | 9,685,297 |
|  |  | Polyclonal Rep 1 | 46,708,750 | 23,354,375 |
|  |  | Polyclonal Rep 2 | 77,858,414 | 38,929,207 |
|  | H3K4me3 – lot comparison | Monoclonal Lot 8 Rep 1 | 83,285,982 | 41,642,991 |
|  |  | Monoclonal Lot 8 Rep 2 | 69,829,160 | 34,914,580 |
|  |  | Monoclonal Lot 9 Rep 1 | 73,635,980 | 36,817,990 |
|  |  | Monoclonal Lot 9 Rep 2 | 68,671,138 | 34,335,569 |
|  | H3K4me3 | Monoclonal Rep 1 | 10,769,082 | 5,384,541 |
|  |  | Monoclonal Rep 2 | 14,706,444 | 7,353,222 |
|  |  | Monoclonal Rep 2 | 14,864,810 | 7,432,405 |
|  |  | Monoclonal Rep 4 | 11,230,992 | 5,615,496 |
|  |  | Polyclonal Rep 1 | 13,184,088 | 6,592,044 |
|  |  | Polyclonal Rep 2 | 10,007,054 | 5,003,527 |
|  |  | Polyclonal Rep 3 | 11,968,378 | 5,984,189 |
|  |  | Polyclonal Rep 4 | 25,205,992 | 12,602,996 |
|  | H3K9me3 | Monoclonal Rep 1 | 75,625,321 | 37,812,661 |
|  |  | Monoclonal Rep 2 | 37,480,051 | 18,740,026 |
|  |  | Monoclonal Rep 2 | 45,075,156 | 22,537,578 |
|  | H3K27ac | Monoclonal Rep 1 | 42,775,420 | 21,387,710 |
|  |  | Monoclonal Rep 2 | 72,589,124 | 36,294,562 |
|  |  | Polyclonal Rep 1 | 60,003,204 | 30,001,602 |
|  |  | Polyclonal Rep 2 | 28,926,788 | 14,463,394 |
|  | H3K27me3 | Monoclonal Rep 1 | 19,964,374 | 9,982,187 |
|  |  | Monoclonal Rep 2 | 24,909,516 | 12,454,758 |
|  |  | Monoclonal Rep 2 | 31,796,524 | 15,898,262 |
|  |  | Monoclonal Rep 4 | 25,983,780 | 12,991,890 |
|  |  | Polyclonal Rep 1 | 22,909,534 | 11,454,767 |
|  |  | Polyclonal Rep 2 | 28,428,020 | 14,214,010 |
|  |  | Polyclonal Rep 3 | 19,843,850 | 9,921,925 |
|  |  | Polyclonal Rep 4 | 37,134,412 | 18,567,206 |
|  | WCE | Rep 1 | 15,726,808 | 7,863,404 |
| mES | H3K4me1 | Monoclonal Rep 1 | 76,330,698 | 38,165,349 |
|  |  | Monoclonal Rep 2 | 98,191,020 | 49,095,510 |
|  | H3K4me3 | Monoclonal Lot 8 Rep 1 | 81,525,630 | 40,762,815 |
|  |  | Monoclonal Lot 8 Rep 2 | 80,975,708 | 40,487,854 |
|  |  | Monoclonal Lot 9 Rep 1 | 72,554,992 | 36,277,496 |
|  |  | Monoclonal Lot 9 Rep 2 | 73,758,624 | 36,879,312 |
|  | H3K27ac | Monoclonal Rep 1 | 161,354,476 | 80,677,238 |
|  |  | Monoclonal Rep 2 | 161,371,680 | 80,685,840 |
|  | H3K27me3 | Monoclonal Rep 1 | 75,925,468 | 37,962,734 |
|  |  | Monoclonal Rep 2 | 164,450,620 | 82,225,310 |

B.
|  |  | Average Reads<br>After<br>Downsampling | Average Pairs<br>After<br>Downsampling | Pairs in Merged<br>Dataset |
| --- | --- | --- | --- | --- |
| K562 | H3K27ac | 25,808,498 | 12,904,249 | 25,808,498 |
|  | H3K27me3 | 18,616,216 | 9,308,108 | 27,924,324 |
|  | H3K4me1 | 35,539,733 | 17,769,867 | 35,539,733 |
|  | H3K4me3 | 9,204,667 | 4,602,334 | 18,409,335 |
|  | H3K9me3 | 28,479,015 | 14,239,507 | 28,479,015 |

